# Epigenetic dynamics of centromeres and neocentromeres in *Cryptococcus deuterogattii*

**DOI:** 10.1101/2020.11.05.369892

**Authors:** Klaas Schotanus, Vikas Yadav, Joseph Heitman

**Affiliations:** Department of Molecular Genetics and Microbiology, Duke University Medical Center, 322 CARL Building, Box 3546, Research Drive, Durham, NC 27710 USA

## Abstract

Deletion of native centromeres in the human fungal pathogen *Cryptococcus deuterogattii* leads to neocentromere formation. Native centromeres span truncated transposable elements, while neocentromeres do not and instead span actively expressed genes. To explore the epigenetic organization of neocentromeres, we analyzed the distribution of the heterochromatic histone modification H3K9me2, 5mC DNA methylation and the euchromatin mark H3K4me2. Native centromeres are enriched for both H3K9me2 and 5mC DNA methylation marks and are devoid of H3K4me2, while neocentromeres do not exhibit any of these features. Neocentromeres in *cen10*Δ mutants are unstable and chromosome-chromosome fusions occur. After chromosome fusion, the neocentromere is inactivated and the native centromere of the chromosome fusion partner remains as the sole, active centromere. In the present study, the active centromere of a fused chromosome was deleted to investigate if epigenetic memory promoted the re-activation of the inactive neocentromere. Our results show that the inactive neocentromere is not re-activated and instead a novel neocentromere forms directly adjacent to the deleted centromere of the fused chromosome. To study the impact of transcription on centromere stability, the actively expressed *URA5* gene was introduced into the CENP-A bound regions of a native centromere. The introduction of the *URA5* gene led to a loss of CENP-A from the native centromere, and a neocentromere formed adjacent to the native centromere location. Remarkably, the inactive, native centromere remained enriched for heterochromatin, yet the integrated gene was expressed and devoid of H3K9me2. A cumulative analysis of multiple CENP-A distribution profiles revealed centromere drift in *C. deuterogattii*, a previously unreported phenomenon in fungi. The CENP-A-binding shifted within the ORF-free regions and showed a possible association with a truncated transposable element. Taken together, our findings reveal that neocentromeres in *C. deuterogattii* are highly unstable and are not marked with an epigenetic memory, distinguishing them from native centromeres.

## Introduction

To undergo proper cell division, chromosomes require a functional centromere that mediates binding of the kinetochore and microtubules to the chromosome [1]. In most organisms, the kinetochore assembles on a specific centromeric histone H3 variant (known as CENP-A, CENH3, or Cse4 depending on the species), which replaces the canonical histone H3 within the centromere [2]. Species lacking CENP-A are either holocentric or differ in their kinetochore structural organization [3–6]. The majority of studied centromeres are regional centromeres that are epigenetically regulated; there are however some exceptions, such as point centromeres that are sequence-dependent [7,8].

Fungal regional centromeres are well studied; in particular, the centromeres of *Candida albicans* and closely related species have been the subject of extensive investigation [9–11]. The centromeres in *C. albicans* are small (ranging from 3.5 to 5 kb) and are flanked by pericentric regions that harbor either inverted repeat or long-terminal repeat sequences, except in the case of chromosome 7 [12–14]. The pericentric regions are enriched for the euchromatic histone marks H3K9Ac and H4K16Ac but are hypomethylated at H3K4, a feature of heterochromatic chromatin [14]. Histone H3 is depleted at the core centromeres of *C. albicans* and H3K9 methylation is absent in the *C. albicans* genome, as this pathway was lost during evolution [15,16]. The mixture of euchromatin and heterochromatin results in reduced expression of genes located in close proximity to the centromeres. In contrast, native centromeres of *Schizosaccharomyces pombe*, another ascomycetous fungus, have a higher-order structure with a CENP-A-enriched central core, flanked by heterochromatic inner and outer repeats that serve as pericentric regions [17]. The pericentric regions are enriched for H3K9me2/3, and this modification is RNAi-dependent and essential for CENP-A localization to the central core [18,19].

The centromeres of basidiomycete *Cryptococcus neoformans* are enriched for both full-length and truncated transposable elements and previous studies have shown that these elements are silenced via RNAi [20–24]. The centromeres of *C. neoformans* are also enriched for heterochromatic marks H3K9me2 and 5mC DNA methylation and in contrast to *S. pombe*, heterochromatin formation in *C. neoformans* is not mediated by small RNAs [22,25,26]. A sister species of *C. neoformans* is *C. deuterogattii* and this is the only member of the *Cryptococcus* pathogenic species complex that is RNAi-deficient [27,28]. The loss of RNAi has been correlated with the presence of only truncated transposable elements within centromeres whose length has been significantly reduced (average length of 14.5 kb) as compared to RNAi-proficient *Cryptococcus* species [17,24]. Despite having only truncated centromeric transposable elements, phylogenetic analysis showed that the transposable elements in *C. deuterogattii* are still similar to those of other members of the species complex [21]. The majority of these truncated centromeric transposable elements are Ty3-Gypsy family retroelements (Tcn1-Tcn5) with one Ty1-Copia element (Tcn6) among them.

Functional studies to analyze and describe *de novo* centromere formation are possible following the deletion of a native centromere. Deletion of centromeres frequently results in neocentromere formation in genomic hotspots, and these are often formed in the vicinity of the native centromere [30]. This has been observed in model organisms such as *C. albicans* and also in chicken DT40 cells, and might be attributable to the presence of a “CENP-A cloud” [31–36]. The CENP-A cloud consists of free, non-incorporated CENP-A molecules that are present in close proximity to the centromeres at a concentration that decreases with increasing distance from the centromeres [36]. It is important to note that centromeres in most fungi are clustered at one focus within the nucleus, which suggests that, the CENP-A cloud might be a specialized part of the nucleus designated for centromeres.

Unlike the native centromeres in *C. albicans*, neocentromeres are not associated with flanking repeats and genes spanned by CENP-A are silenced [34]. Interestingly, all neocentromeres of chromosome 7 were found to form next to the native centromere, while only half of the neocentromeres on chromosome 5 formed in close proximity to the native centromere [33,34]. The lengths of neocentromeres varied from 2 to 7 kb, suggesting neocentromeres in this species can be flexible, unlike its native centromeres. While the epigenetic properties of neocentromeres are not yet defined in this species, the formation of a neocentromere changes the genome-wide interaction landscape [35]. A recent study also showed that neocentromeres in *C. albicans* are confined to the pericentric region that is defined by cis- and trans-interactions of the chromatin [37].

Similar to *C. albicans*, the majority (76%) of neocentromeres in chicken DT40 cells form in close proximity to the native centromere [32]. Chickens have a diploid genome, but the Z sex chromosome is present in only one copy in DT40 cells (ZW/female cells). Both the native centromere and neocentromeres of chromosome Z lack repeats, do not contain a specific DNA motif and are similar in length (∼35 to 45 kb). Based on chromatin immunoprecipitation sequencing (ChIP-seq) and immunofluorescence microscopy, native centromeres of chromosomes 5, 27, and Z, as well as all neocentromeres, are not enriched for H3K9me3. This is in contrast to the native centromeres of chromosomes 1 and 2, which span repeats and are enriched for H3K9me3. Despite the lack of H3K9me3, the expression level of a gene (*MAMDC2*) within a neocentromere is reduced by 20-to 100-fold relative to wild-type expression levels that was also reflected by a reduction of H3K4me2 levels [32]. These findings suggest heterochromatin is not essential for centromere function in DT40 cells and that heterochromatin may serve to repress only repetitive regions.

Neocentromeres in *S. pombe* form adjacent to heterochromatic regions, close to the telomeres, where pre-existing repeats mimic the higher-order repeat architecture of native centromeres [38]. Similar to *C. albicans, S. pombe* neocentromeres can span genes that are silenced due to CENP-A binding [38]. Interestingly, deletion of native centromeres in wild-type *S. pombe* leads to either neocentromere formation or chromosome fusion, while centromere deletion in H3K9me2-deficient cells results in only chromosome fusion, suggesting that heterochromatin is essential for neocentromere formation and function in *S. pombe* [38,39].

Previously we showed that deletion of a native centromere in *C. deuterogattii* leads to neocentromere formation [40]. Native centromeres and neocentromere are organized in dramatically different fashions. Native centromeres span truncated transposable elements and are located in ORF-free regions. In contrast, neocentromeres of both chromosomes 9 and 10 span genes that continue to be expressed at wild-type levels. The neocentromeres are shorter in length and lack transposable elements. The wild-type expression of neocentromeric genes suggests neocentromeres may not be heterochromatic in *C. deuterogattii*. However, neocentromeres of chromosome 10 were unstable and often led to chromosome-chromosome fusions between sub-telomeric regions. After chromosome fusion, the neocentromeres were inactivated and only the native centromere of the fusion partner chromosome served as the centromere of the fused chromosome [40].

In the current study, we show that deletion of the native centromere in fused chromosomes results in neocentromere formation in close proximity to the native centromere, not at the previous neocentromere location. We find that, in contrast to native centromeres, neocentromeres are not enriched for the heterochromatin mark H3K9me2 or 5mC DNA methylation and maintain their H3K4me2 status. Additionally, integration of a selectable marker gene into a native centromere resulted in a loss of CENP-A binding at the native centromere and the formation of a neocentromere adjacent to the site of the original native centromere. Overall, our results demonstrate that *C. deuterogattii* neocentromeres 1) lack epigenetic marks that are present at native centromeres, and 2) do not retain any epigenetic memory following inactivation.

## Results

### Native centromeres are enriched for the heterochromatic mark H3K9me2 in *C. deuterogattii*

The centromeric regions of the closely related sister species *C. neoformans* are enriched for active transposable elements, small-RNAs, 5mC DNA methylation, and heterochromatin [21,25,26]. In contrast, the sister species *C. deuterogattii* has lost the RNAi pathway and the centromeres lack active transposable elements and contain only truncated remnants of Tcn1– Tcn6 transposable elements located in non-coding, ORF-free chromosomal regions [21]. Except for Tcn6 (Ty1-Copia family), the rest of these Tcn elements are Ty3-Gypsy transposable elements and almost all of these truncated retro-transposable elements are centromeric [21].

As a result, it is possible that transposable element silencing pathways, in addition to RNAi, such as heterochromatin and DNA methylation, could also be redundant and might have been lost or evolved differently in this species. To test if heterochromatin is still functional in *C. deuterogattii*, we performed ChIP followed by high-throughput sequencing (ChIP-seq) with an antibody specific to H3K9me2.

Despite the lack of active transposable elements, the native centromeres were found to be enriched for H3K9me2 and this chromatin mark spanned the entire ORF-free region (Figure 1). Further analysis revealed that the CENP-A-bound region was embedded within the H3K9me2-enriched region and displayed a similar pattern as was previously shown in *C. neoformans* (Figure S1) [25]. Telomeres and sub-telomeric regions were also found to be enriched for H3K9me2. In addition to the expected regions, we identified several short chromosomal regions (non-telomeric and non-centromeric regions) that were enriched for H3K9me2; however, compared to the centromeres, the level of H3K9me2 enrichment was at least five-fold lower in these regions (Table S3). Some of these H3K9me2-enriched regions spanned genes whose products are predicted to be involved in sugar metabolism or drug transport.

**Figure 1.**
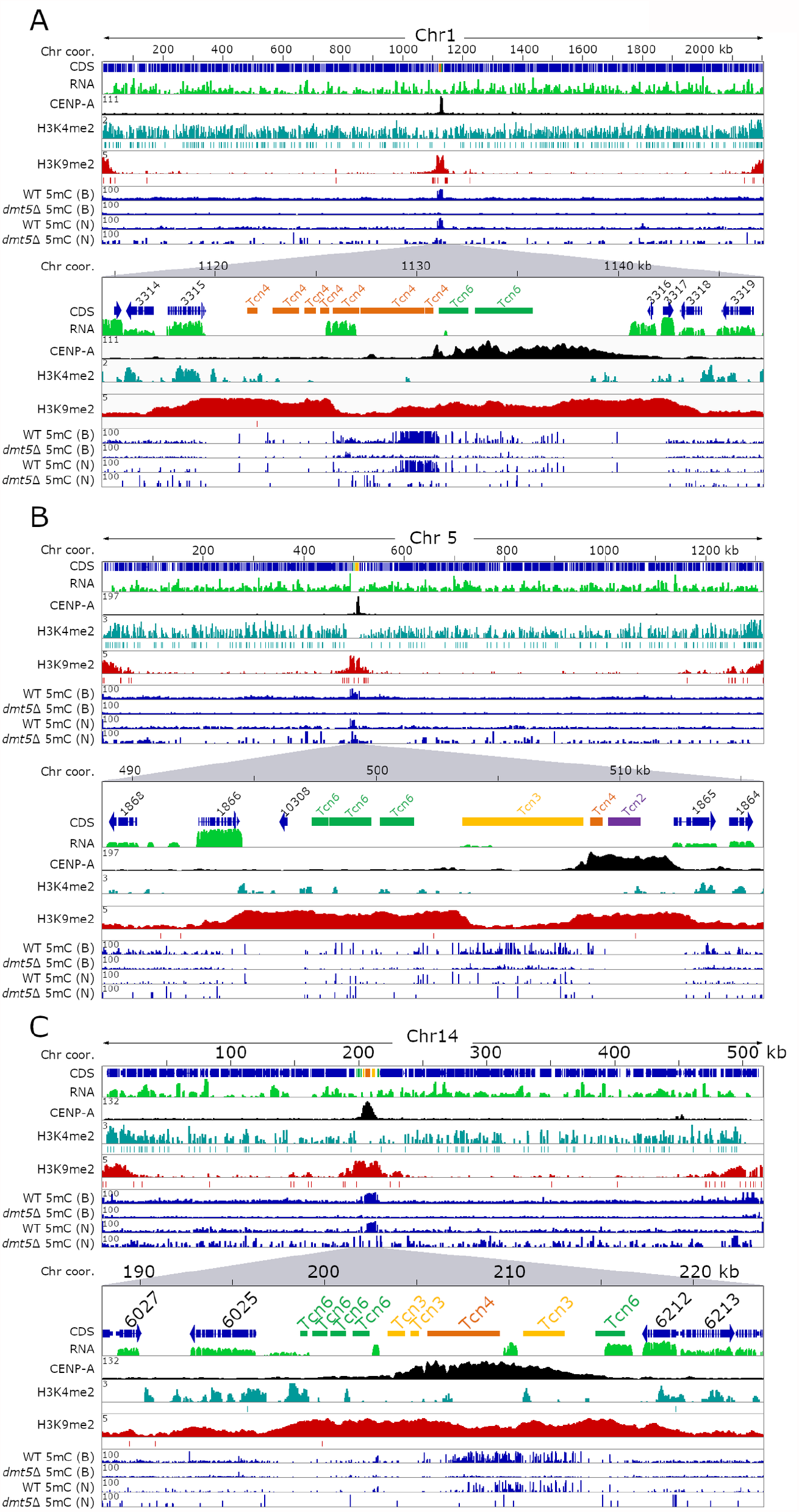
Heterochromatin marks H3K9me2 and 5mC DNA methylation do not always coincide on *C. deuterogattii* centromeres. For each panel, the full-length chromosome (Chr) is on the top and the lower section shows a region spanning the native centromere with 2-4 flanking genes. Each panel shows the chromosome coordinates, with arrows indicating the gene coding sequences (CDS), TEs present in centromeres, RNA-seq shown in green, CENP-A reads indicating the native centromere shown in black, H3K4me2 enrichment is in turquoise, H3K9me2 enrichment is in red, and 5mC enrichment is shown in blue (for both the wild type and *dmt5*Δ). 5mC data is derived from either bisulfite sequencing (B) or Nanopore sequencing (N). The enrichment of each sample over the input signal is shown on the left of each panel. MACS2 was used to validate significant peaks and indicated with a line below the whole peak. Panel **(A)** shows Chr1, panel **(B)** shows Chr5, and panel **(C)** shows Chr14.

In addition to analysis of the heterochromatic mark, we performed ChIP-seq with an antibody against the euchromatic histone mark H3K4me2. This histone mark colocalizes with actively expressed genes and was mostly absent from centromeres (Figure 1). This binding pattern also reflected the poor transcription observed at centromeres in this species. Combined together, these results revealed that centromeres in *C. deuterogattii* are heterochromatic in nature despite the loss of RNAi and the absence of full-length, active transposons.

### Correlation between 5mC DNA methylation and H3K9me2 across centromeres

We previously reported that the DNA methyltransferase gene *DMT5* was truncated in the *C. deuterogattii* R265 reference genome, and 5mC methylation was thought to be entirely absent based on PCR-based assays with methylation-sensitive restriction enzymes [21]. An updated gene annotation of the R265 reference genome and re-mapping of RNA-seq data revealed that the *DMT5* gene was previously mis-annotated and it could encode a putative fully functional protein (Figure S2) [41]. To analyze the function of 5mC DNA methylation, the *DMT5* gene was deleted by homologous recombination and bisulfite sequencing was performed with DNA isolated from the wild-type strain and a *dmt5*Δ deletion mutant (Figure S3A). Two independent DNA methylation analysis methods, bisulfite sequencing, and DNA methylation enrichment from nanopore sequencing showed that centromeres are enriched for 5mC in *C. deuterogattii*. However, methylation levels were significantly reduced compared to the *C. neoformans* wild-type reference strain H99 (Figure 1 and S1). Furthermore, the 5mC was observed only in a subset of centromeres, and even in these cases, it was localized to specific regions instead of pan-centromere, unlike *C. neoformans*. As expected, 5mC levels were abolished in the R265 *dmt5*Δ mutant strain. The bisulfite sequencing analysis also confirmed that 5mC was lacking in the R265 genomic regions that were previously analyzed by PCR following methylation-sensitive restriction enzyme digestion [21].

5mC DNA methylation in *C. neoformans* co-localized with H3K9me2 and DNA methylation was also found to be dependent on the presence of H3K9me2 [26]. In contrast to *C. neoformans*, our results show that H3K9me2 and 5mC are not well-correlated with each other in *C. deuterogattii*. For example, the two longest members of the retro-transposable elements family, Tcn3 (∼5 kb), located in *CEN3* and *CEN5*, lack both CENP-A and H3K9me2 enrichment but are among the most abundant 5mC-enriched regions in the *C. deuterogattii* genome (Figure 1). The remaining four shorter Tcn3 elements (∼0.4 to 2.2 kb) are enriched for H3K9me2 and CENP-A but not significantly enriched for 5mC. Transposable elements Tcn4 and Tcn6 are also enriched with both 5mC and H3K9me2 suggesting that the two modifications can occur at the same locus. Furthermore, *CEN4, CEN6*, and *CEN7* completely lack 5mC enrichment but are significantly enriched for H3K9me2 (Figure S1). Based on these results, we conclude that even though 5mC methylation is present in *C. deuterogattii*, the overall level is significantly reduced compared to *C. neoformans*. In addition, two heterochromatic marks, 5mC and H3K9me2, differ significantly in terms of their binding between the two species.

### Neocentromeres are not enriched for the heterochromatic mark H3K9me2 or 5mC

Previously, we deleted native *CEN10* and obtained multiple isolates in which neocentromeres were formed [40]. These neocentromeres span actively-expressed genes and the majority of the neocentromeres are located in close proximity to the deleted, native centromere [40]. To test if there is a correlation between histone marks and the chromosomal location of neocentromeres, we analyzed the chromatin of wild-type chromosome 10 (Figure 2A). As expected, euchromatin (H3K4me2) is spread along the length of the chromosome and co-localizes with the genes, contrasting with the heterochromatin mark (H3K9me2) that is enriched at the subtelomeric regions and the centromere.

**Figure 2.**
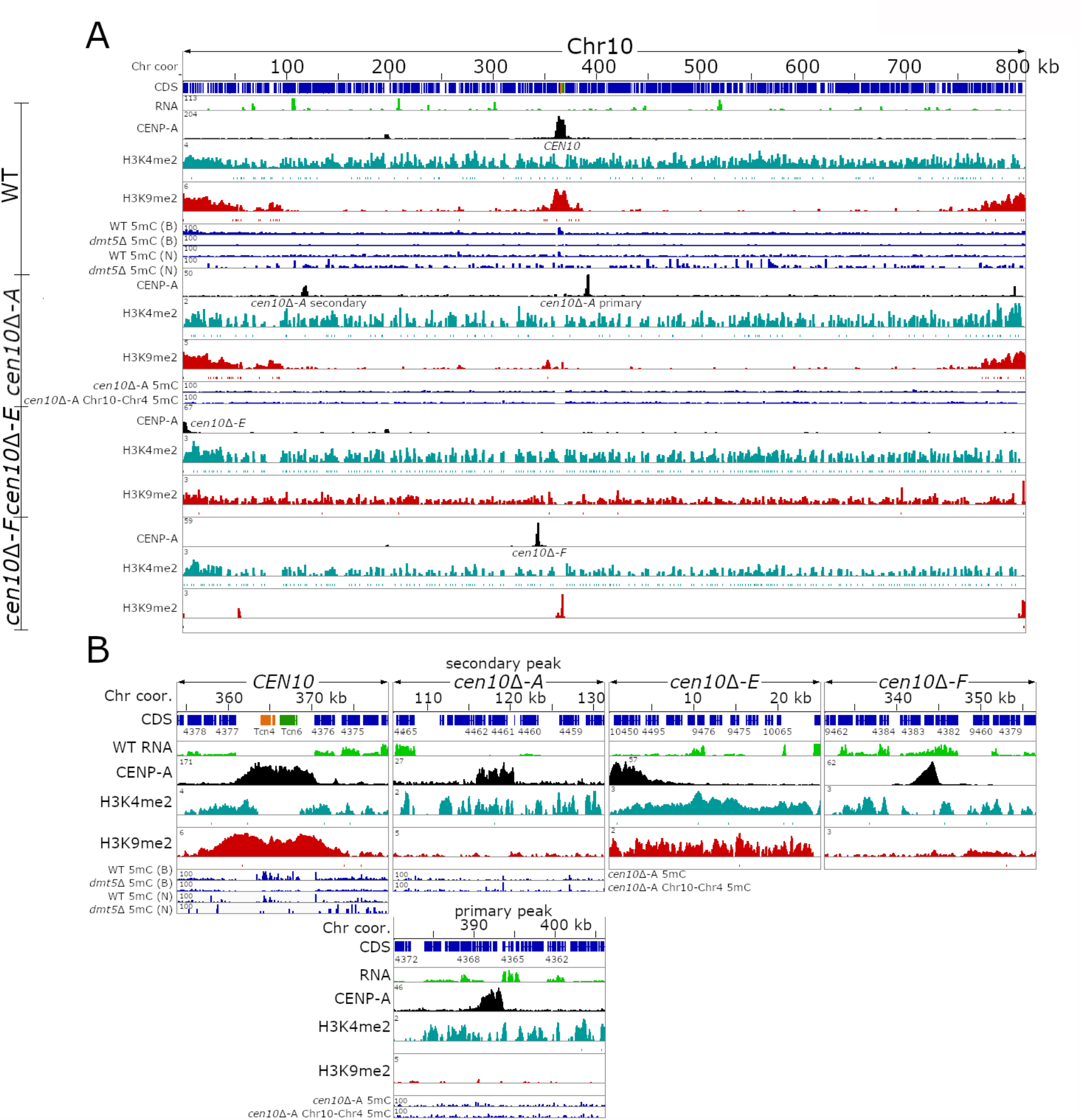
*C. deuterogattii* neocentromeres are not enriched for H3K9me2. For each panel, the chromosome coordinates are indicated. CDS are shown in blue, TEs are presented as per the specific TE family color, and RNA-seq is shown in green. CENP-A ChIP-seq is shown in black, ChIP-seq for H3K4me2 is shown in turquoise, and ChIP-seq for H3K9me2 is shown in red. DNA methylation for the wild-type is shown in blue and the source of the 5mC data is indicated by B (Bisulfite sequencing) or N (Nanopore sequencing). **(A)** The top panel shows the distribution of epigenetics marks of the full-length wild-type Chr10. The lower panels show the epigenetic marks of the full-length Chr10 of *cen10*Δ-*A, cen10*Δ-*E*, and *cen10*Δ-*F* strains. As shown *cen10*Δ-*A* has two neocentromeric peaks, while the other isolates only have one centromere on chromosome 10. For this isolate, DNA methylation enrichment for *cen10*Δ-*A* is shown and in addition the 5mC enrichment for the isolate with the chr10-4 fusion resulting in the inactivation of the neocentromere is shown for comparison. **(B)** Detailed centromeric view of the centromere or neocentromeres of Chr10 of the wild-type, *cen10*Δ-*A, cen10*Δ-*E*, and *cen10*Δ-*F*.

To test if *cen10*Δ neocentromeres are enriched for H3K4me2 and H3K9me2, ChIP-seq with antibodies specific for either H3K4me2 or H3K9me2 were performed for three *cen10*Δ strains containing neocentromeres (Figure 2A, 2B). Isolate *cen10*Δ-*A* has two CENP-A-enriched regions of which one is the primary peak and the second is a less abundantly enriched peak [40]. Isolate *cen10*Δ-*E* has a neocentromere directly flanking the left telomere and isolate *cen10*Δ-*F* has a neocentromere proximal to the deleted native centromere [40]. After mapping the H3K4me2 and H3K9me2 ChIP-seq reads to the genome, we observed that the neocentromeres of *cen10*Δ-*A* and *cen10*Δ-*F* lacked H3K9me2 enrichment (Figure 2A).

The subtelomeric regions of all 14 chromosomes as well as the native centromeres of the other 13 chromosomes were all still enriched for H3K9me2 as expected. As the neocentromere of *cen10*Δ-*E* is telocentric, it was enriched with H3K9me2. However, because the sub-telomeric regions are already modestly enriched for H3K9me2 in the wild type, we think that the enrichment in this isolate simply reflects that the neocentromere is located in a sub-telomeric region that was already modified. Overall, our results show that H3K9me2 is not required for neocentromere function in *C. deuterogattii*. Furthermore, all *cen10*Δ neocentromeres were found to be enriched for H3K4me2, and the enrichment levels for the neocentromeric chromosomal locations are similar in *cen10*Δ mutants and wild-type (Figure 2B). These results suggest that neocentromere locations are not modified with respect to their H3K9me2 or H3K4me2 marks.

We also tested for 5mC methylation at the neocentromere locations and performed Oxford Nanopore sequencing for four *cen10*Δ mutants (Figure 2, Figure S4). As shown in Figure 1 and Figure 2, the nanopore sequencing data provides an accurate estimation of 5mC and thus was used to estimate the 5mC DNA methylation of these neocentromeric isolates. All of the neocentromere regions analyzed lacked any enrichment for 5mC DNA methylation in the wild-type and as well as in neocentromere-harboring strains (Figure S4). This is not surprising provided that all pathogenic *Cryptococcus* species were shown to harbor only the maintenance DNA methyltransferase Dnmt5 and have lost the *de novo* DNA methyltransferase enzyme [26]. In summary, native centromeres are located in a large ORF-free region, enriched for truncated transposable elements, H3K9me3, and 5mC, whereas neocentromeres are smaller, span actively expressed genes, and lack enrichment for H3K9me2 or 5mC (Figure S5).

### Inactive neocentromeres lack epigenetic memory

In a previous study, we observed that *cen10*Δ mutant isolates with a neocentromere exhibited aneuploidy for chromosome 10 and produced both large and small colonies. Isolation and analysis of larger colonies formed at 37°C revealed that chromosome-chromosome fusions had occurred in which chromosome 10 was fused with another chromosome [40]. After chromosome-chromosome fusion, the neocentromere was inactivated and the native centromere of the fusion partner chromosome functioned as the sole, active centromere. As the neocentromere was inactivated in the fused chromosome, centromeric proteins were displaced from the neocentromere locus, and we experimentally confirmed this by showing that the inactivated neocentromere lacked CENP-A. However, we hypothesized that the inactive neocentromere might still bear some unidentified epigenetic marks, and if so, these marks might render this a preferred site for subsequent neocentromere formation.

To investigate whether such epigenetic memory existed at the inactive neocentromeric location, we deleted the active centromere of the fused chromosome and performed ChIP-seq analysis for CENP-A (Figure 3). In the previous study, we described three chromosome fusions in more detail and for this experiment, we focused on an isolate (KS123) in which chromosome 10 was fused with chromosome 4 [40]. This strain is derived from a dineocentric *cen10*Δ isolate with a primary CENP-A peak located in close proximity to the deleted native *CEN10* and a smaller secondary CENP-A peak with reduced CENP-A levels located 116 kb from the 5’ telomere (Figure 3B). After the chromosome-chromosome fusion event, the neocentromeres were inactivated and the native centromere of the fusion partner (in this example chromosome 4) became the active centromere for the chromosome fusion. We hypothesize that the dineocentric nature of the parental isolate could increase the chances of detecting re-activation of inactive neocentromeric locations upon deletion of *CEN4* in this strain. Thus, by using CRISPR-Cas9, we deleted native *CEN4* and obtained two independent *cen10*Δ *cen4*Δ mutants (Figure S3B). In addition, we also deleted *CEN4* in the wild type (WT) and obtained two independent *cen4*Δ mutants to serve as controls. All four mutants lacking *CEN4* were subjected to CENP-A ChIP-seq analysis and reads were mapped to the genome assembly.

**Figure 3.**
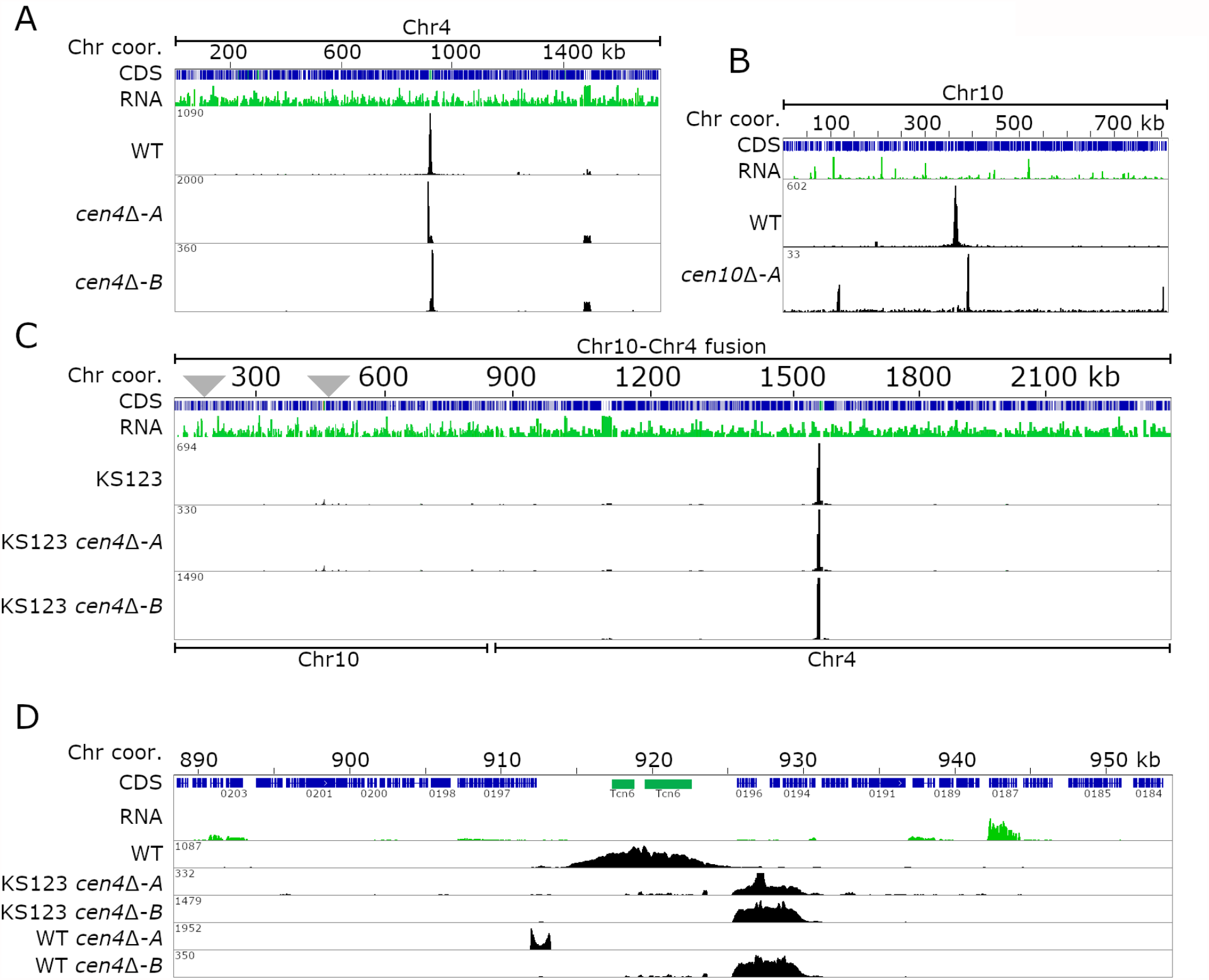
Neocentromeres in *C. deuterogattii* lack epigenetic memory. For each panel, the chromosome is indicated on top of the panel, followed by the chromosome coordinates. The genes (CDS) are shown in blue and the RNA-seq is shown in green. The CENP-A tracks are shown in black. **(A)** Full-length chromosomal view for Chr4, showing the position of *CEN4* and the two neocentromeric strains obtained in the wild-type karyotype. **(B)** The panel shows the previously obtained parental dineocentromeric *cen10*Δ-*A* strain. **(C)** A full-length chromosomal view of two *cen4*Δ mutants in a Chr10-Chr4 fusion derived from the *cen10*Δ mutant background (KS123) is shown. As shown in panel B, the initial *cen10*Δ-*A* mutant had two CENP-A-enriched neocentromeric regions, and these inactivated neocentromeres are indicated with gray triangles. **(D)** Detailed view of native *CEN4* and neocentromeric regions. As shown with two independent *cen4*Δ deletions isolated in the strain harboring the fused chromosome, the neocentromere was inactivated after chromosome fusion. The panel with *CEN4* shows that neocentromeres formed proximal to the native *CEN4*, and this was independent of the presence of the inactive neocentromere. In addition to *CEN4* deletion in the fused strain, two independent *cen4*Δ in wild-type strains show neocentromere formation in close proximity of the native centromere, and one of these neocentromere co-localizes with neocentromeres in the fused strain.

Interestingly, in all four mutants, the neocentromeres formed close to the location of native *CEN4* (Figure 3A, C and D). In three cases, neocentromeres formed in the same chromosomal location, directly flanking the native centromere on the 3’ side. These neocentromeres were smaller in size (∼5 kb) than the native *CEN4* (∼10 kb) and spanned three genes (CNBG_0196, CNBG_0195, and CNBG_0194). CNBG_0196 encodes a serine/threonine-protein kinase, CNBG_0195 encodes a purine-specific oxidized base lesion DNA N-glycosylase, and CNBG_0194 encodes a protein predicted to localize to the endoplasmic reticulum. In the fourth case, the neocentromere formed next to the native centromere on the 5’ side. This neocentromere was much smaller in size (∼1.3 kb) than the native *CEN4* as well as the other three neocentromeres and spanned only the first two exons of the CNBG_0197 gene, which encodes topoisomerase II alpha-4. Unlike *cen10*Δ original mutants, *cen4*Δ mutants in the otherwise wild-type background had uniform colony sizes, as was previously observed with *cen9*Δ mutants, suggesting a stable neocentromere [40]. Taken together, these results provide evidence that inactivated neocentromeres in *C. deuterogattii* do not possess any epigenetic memory and suggest that *de novo* neocentromere formation may instead occur due to the presence of a “CENP-A cloud”.

### Chromosome fusion leads to centromere inactivation instead of a dicentric chromosome

As mentioned earlier, the parent of strain KS123 was a *cen10*Δ isolate with two neocentromeres. This prompted us to test whether a chromosome with two native centromeres would result in a stable dicentric chromosome in *C. deuterogattii*. To study this, we used CRISPR-Cas9 and generated a fusion of chromosomes 4 and 10. Using this approach, two isolates were obtained in which the targeted chromosome fusion was confirmed based on Pulsed-field gel electrophoresis (PFGE) as indicated by the absence of bands corresponding to chromosomes 4 and 10 (Figure S6). However, from the EtBr stained banding pattern, it appears that the dicentric chromosome in one of the transformants (dicentric isolate 1) broke resulting in a new chromosomal band at ∼890 kb (Figure S6C and D). The second transformant (dicentric isolate 2) did not show any new band suggesting that either the dicentric chromosome was stable or the PFGE was not able to resolve the new chromosomes. To confirm the chromosome fusion, nanopore sequencing was conducted for both putative dicentric isolates. For dicentric isolate 1, a scaffold (443 kb) with chromosome 10 fused to a 60 kb region of chromosome 4 was identified. Interestingly, the same 60 kb region appeared duplicated in the genome and was also present on a scaffold (386 kb) that corresponds to a part of chromosome 4 (Figure S7). Short-read Illumina sequencing confirmed that the 60 kb region was indeed duplicated. We think that, in this isolate, the dicentric chromosome broke, and two new chromosomal ends were generated along with a 60 kb segmental duplication shared by the new versions of chromosomes 4 and 10.

For dicentric isolate 2, nanopore sequencing suggested the presence of a stable chromosome 4-10 fusion. To test if either centromere 4 or 10 was inactivated after chromosome fusion in this isolate, ChIP-qPCR analyses targeting CENP-A enrichment of *CEN4* and *CEN10* of the putative dicentric chromosome were performed (Figure S6E). Compared to the positive control (*CEN6*), the CENP-A enrichment for *CEN4* was 2-fold higher, while the CENP-A enrichment for *CEN10* was 0.5-fold lower. This result suggests that the majority of cells in the population have an active *CEN4* whereas a smaller proportion of cells have *CEN10* as the active centromere. However, we cannot rule out the possibility that both centromeres might be active in some cells and still be stable.

This small-scale experiment shows that following chromosome-chromosome fusion, in one example the resulting dicentric chromosome was broken prior to analysis of the fusion in detail, suggesting that a dicentric chromosome is unstable in *C. deuterogattii*. In contrast, the second chromosome fusion analyzed was found to be stable although we observed differential loading of CENP-A at the two centromeric regions suggesting that this mutant may not represent a true dicentric, but rather that one or the other centromere was active in different cells in the population. This second pattern might be similar to that observed for two earlier described *cen10*Δ mutants that harbor both a primary and a secondary CENP-A peak. However, we note that this hypothesis was only tested with two biological replicates and further studies will be necessary to address the relative frequency of these pathways for resolving the fate of unstable dicentric chromosomes.

### Integration of a *URA5* transgene into *CEN2* results in neocentromere formation

*C. deuterogattii* neocentromeres are present in gene-rich regions and span genes that are actively expressed despite CENP-A binding. This prompted us to test the impact of an insertion of an actively transcribed gene into a native centromere might have on centromeric chromatin. To test this, we introduced the *URA5* gene into the CENP-A-binding region of *CEN2* by homologous recombination (Figure 4). *CEN2* was selected because the CENP-A enriched region for *CEN2* spans a relatively small number of truncated TEs and other simple repeats, which allowed us to design a *CEN2*-specific homologous recombination insertion product that we hypothesized would more facilely integrate into this less repetitive centromere compared to other more repetitive centromeres [21]. We hypothesized five possible outcomes from this experiment: 1) CENP-A would cover the *URA5* gene, making the CENP-A-bound region larger than the native centromere; 2) the *URA5* gene might divide the CENP-A enrichment into two independent regions; 3&4) only one of the regions (either left or right) flanking *URA5* would be enriched for CENP-A, generating a smaller centromere; or 5) *URA5* integration might abolish *CEN2* function leading to neocentromere formation (Figure S8).

**Figure 4.**
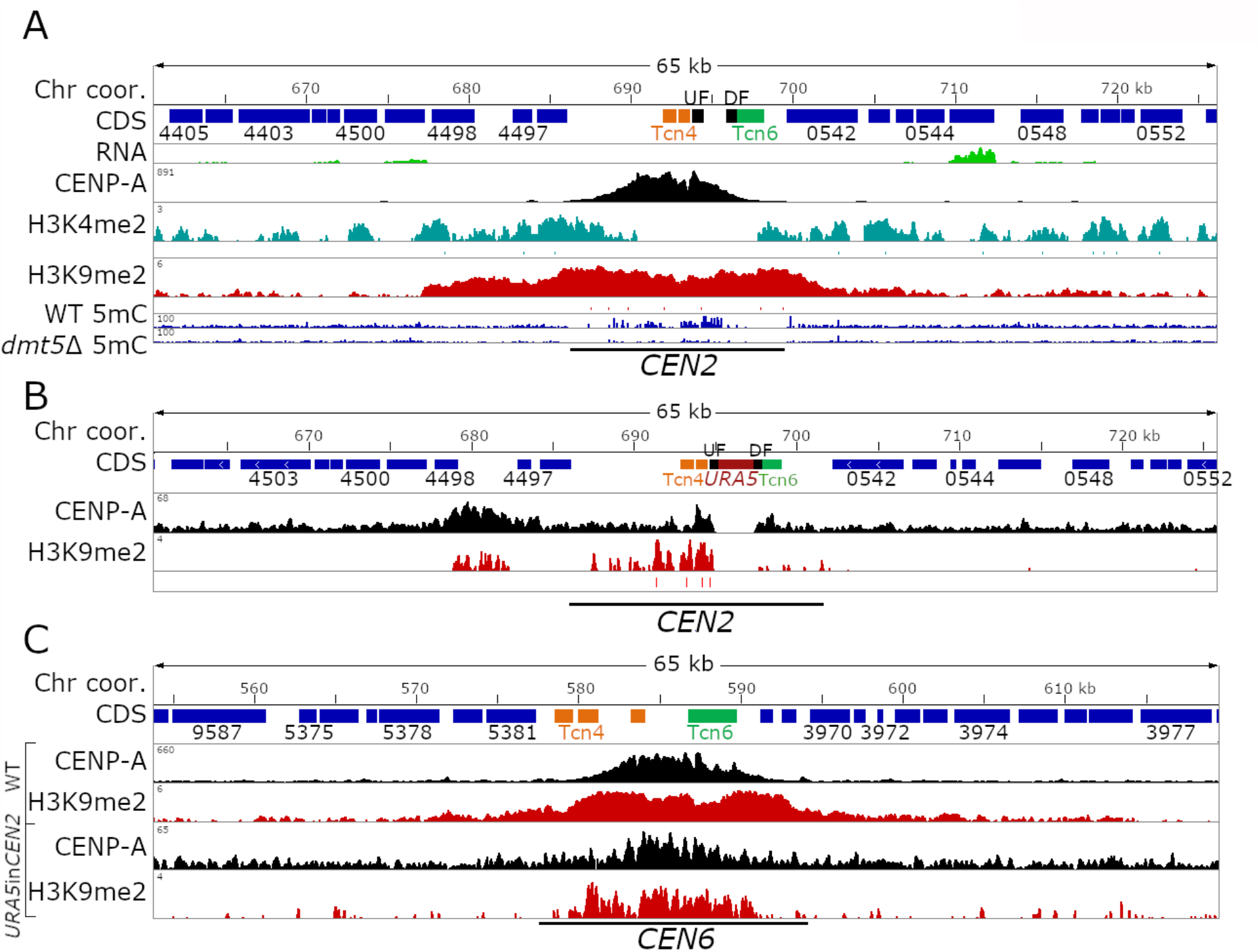
Insertion of a transgene into a native centromere shifts the CENP-A binding. For all three panels, a detailed centromeric view (65kb) of *CEN2* or *CEN6* is shown. Chromosome coordinates are shown on top, CDS are shown in blue, and truncated transposable elements are indicated with their family color. ChIP-seq is shown in black (CENP-A), turquoise (H3K4me2, only wild-type), and red (H3K9me2). UF and DF indicate the 5’ and 3’ region used for homologous recombination. **(A)** Detailed view of native centromere 2. In addition to the ChIP-seq data, bisulfite sequencing data to analyze 5mC enrichment is shown. **(B)** Detailed view showing CENP-A and H3K9me2 enrichment in the *URA5* insertion strain. The *URA5* gene is shown in red. **(C)** Enrichment level across *CEN6* are shown as control for both ChIP-seq experiments which revealed that one experiment showed better enrichment than the other, albeit in the same regions.

Through homologous recombination, the *URA5* gene was introduced into the CENP-A-enriched region of *CEN2* in an R265 *ura5-*strain. It is important to mention that the *ura5-*strain used harbors a point mutation in the *URA5* gene and not a gene deletion. Following transformation, isolates were selected for growth on SD medium lacking uracil (SD-uracil). Isolates were screened by PCR and one transformant with the desired integration of the *URA5* gene into *CEN2* (*URA5*in*CEN2*) was obtained. Southern blot analysis with a probe corresponding to the *URA5* gene hybridized to only two locations in the genome corresponding to the *ura5*-gene at its native location and the *URA5* gene integrated into *CEN2* confirming a single insertion event of the *URA5* gene in the genome at the desired location in *CEN2* (Figure S9). Growth assays revealed that the *URA5* gene introduced into *CEN2* was expressed since the *URA5*in*CEN2* strain grew on SD-uracil media but not on the media containing the toxin 5-fluoro-orotic acid (5-FOA) (Figure S8B). We further tested the expression of *URA5* by more robust selection assays. For this purpose, wild-type and *URA5*in*CEN2* strains were grown in SD uracil and YPD plates as selective and non-selective conditions. From both these conditions, a large number of cells were then plated on 5-FOA and YPD media in order to determine the population of cells where *URA5* might be silenced. However, both wild-type and *URA5*in*CEN2* strains exhibited similar levels of 5-FOA resistant colonies suggesting that *URA5* in the *URA5*in*CEN2* strain is not silenced at all and behaves in the same fashion as *URA5* in the wild-type strain (Figure S8C).

Subsequently, we performed CENP-A ChIP-seq of the *URA5*in*CEN*2 isolate with cells of this mutant grown on rich YPD media to investigate which of the five hypotheses might be correct. Growth in YPD does not select for cells expressing *URA5* neither does it kill those cells, thus allowing assessment of binding of CENP-A in the absence of external factors. After mapping the ChIP-seq reads to the wild-type reference genome, we found that the CENP-A distribution shifted dramatically and co-localized with a region modestly enriched for H3K9me2, flanking one side of the native *CEN2* in accord with hypothesis 5 i.e. *URA5* integration abolishes *CEN2* function and leads to neocentromere formation (Figure 4 and S8A). Despite the formation of a *de novo* CENP-A peak flanking *CEN2*, there was still a small amount of residual CENP-A binding at the native location. The neocentromere spanned ∼6 kb and included three genes: CNBG_4498 which encodes a pyridoxal kinase, CNBG_10170 which encodes the DNA mismatch repair protein Msh4, and CNBG_4497 which encodes a hypothetical protein.

The shift in CENP-A binding after *URA5* insertion into *CEN2* provided a unique opportunity to test if H3K9me2 enrichment remained at the native location. To characterize H3K9me2 enrichment in this strain (KS174), we performed ChIP-seq and quantified the H3K9me2 enrichment within *CEN2*. This ChIP-seq experiment, however, did not show same level of H3K9me2 enrichment as for the wild-type and as a result all centromeres had lower levels of H3K9me2 enrichment (Figure 4). Nonetheless, the inactivated *CEN2* still has H3K9me2 enrichment, both in transposon-rich and unique regions, even though some regions lacked the modification entirely (Figure 4A and B). Interestingly, the neocentromere region was also found to be enriched for H3K9me2 suggesting that CENP-A binding had no effect on H3K9me2 enrichment. We specifically looked at the H3K9me2 enrichment at the *URA5* gene because the gene is actively expressed. As expected, the *URA5* gene completely lacked the H3K9me2. Combined together, these results suggest that insertion of a transgene into a centromere can induce neocentromere formation. Furthermore, the insertion of the transgene does not have a significant impact on the H3K9me2 enrichment throughout the centromere but only perturbs it locally where the transgene is introduced.

### CENP-A binding drifts within the ORF free region of *CEN5* and *CEN14*

The length of a centromere in *Cryptococcus* species is defined by the distance between the two closest flanking genes [21]. The ORF-free chromosomal region between these two flanking genes is enriched for full-length or truncated transposable elements. Here we showed that most *C. deuterogattii* centromeres have the entire centromeric region enriched with heterochromatin whereas the extent of the CENP-A enrichment represents only a fraction of the total centromere length (Figure 1). Previously it was shown that the CENP-A binding can “move” within the pericentric region and this is called centromeric drift, or shift [42]. For example the CENP-A enriched region drifts within the centromeric region of chromosome 5 and 8 in Maize and closely related species and a similar event was shown to occur in *CEN Z* of DT40 cells [42,43].

To identify the chromosomal locations of the neocentromeric mutants for both this current study and our previous study, we submitted a large number of sequencing reactions for ChIPs with an antibody against CENP-A [40]. In addition to the ChIP-seq, genomes of several centromere mutants were sequenced with Illumina technology and *de novo* genome assemblies showed that the genomes of these mutants were syntenic with the reference genome and no genome rearrangements had occurred [40]. Comparison of the CENP-A profiles of *cen4*Δ, *cen9*Δ, *cen10*Δ, and wild-type strains revealed that the CENP-A-bound regions of most centromeres are co-linear and coincident among the strains sequenced [21,40]. However, for *CEN5* and *CEN14*, CENP-A binding was observed to drift within the ORF-free, heterochromatin-enriched centromeric regions (Figure 5). We also re-mapped the earlier obtained CENP-C ChIP-seq data and this analysis further supported the presence of centromeric drift in the pericentric region [21]. For *CEN5*, centromeric drift was observed to be strongly associated with the presence of a Tcn3 element in the middle of the centromere. None of the strains analyzed in ChIP-seq experiments exhibited CENP-A binding to this Tcn3 element, and instead, the binding was localized to either the 5’ or 3’ of the Tcn3 element. Interestingly this Tcn3 element was also not enriched for H3K9me2 but was enriched for 5mC DNA methylation (Figure 5A and Table S4). There is only one other long Tcn3 element that is located in *CEN3*, and this is also not enriched for CENP-A and H3K9me2. However, CENP-A drift was not observed for *CEN3*, probably because this element is located close (∼1.15 kb) to the 3’ flanking gene of the centromere restricting the CENP-A binding dynamics.

**Figure 5.**
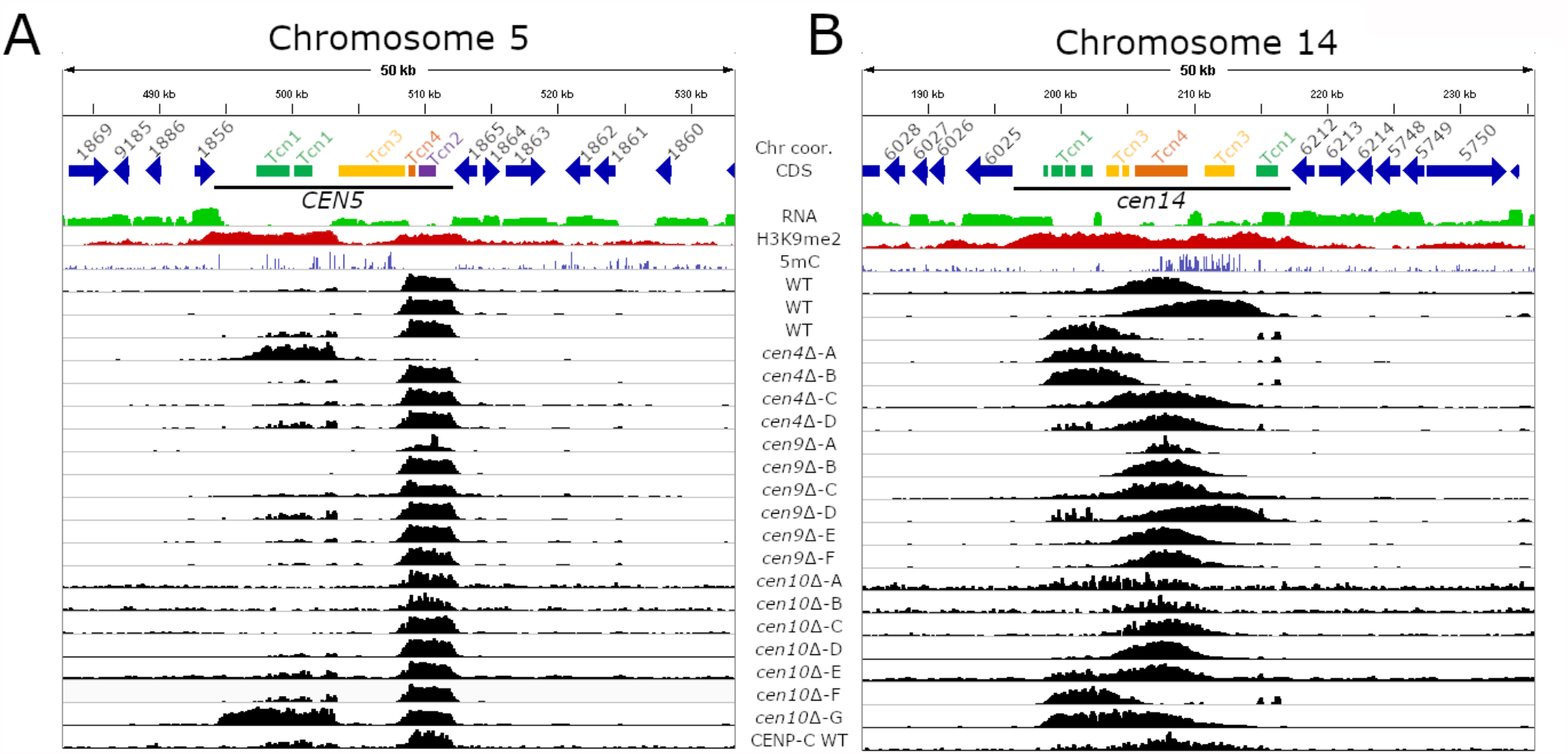
Centromeric drift occurs within ORF-free regions of *C. deuterogattii*. The top panels of both figures show the chromosomal coordinates, genome annotation, RNA-seq, H3K9me2 ChIP-seq, and 5mC for wild-type. CENP-A ChIP-seq is shown in black and the final panel shows CENP-C ChIP-seq. **(A)** Centromeric drift in *CEN5* is correlated with the presence of a large, but truncated Tcn3 element. This truncated Tcn3 element functions as a barrier and for all isolates with the exception of *cen10*Δ-G, CENP-A is present on only one side of the truncated Tcn3 element. **(B)** Centromere drift occurs in *CEN14* and the CENP-A peaks drift within the ORF-free region of *CEN14* and lack a defining feature.

The centromeric drift in *CEN14* was less distinct and the CENP-A-enriched region shifted within the pericentric region and no definitive barrier could be identified (Figure 5B). However, *CEN14* also harbors 3 out of 4 smaller copies of Tcn3, indicating that there might be some relation of Tcn3 transposon with CENP-A drift observed in this centromere. Overall, these results reveal the presence of centromere drift in *C. deuterogattii*, a phenomenon that is not yet reported in fungi. Further studies will shed more light on this phenomenon and its correlation with Tcn3 or other retrotransposons in other species.

## Discussion

### Native centromeres, but not neocentromeres, are heterochromatic in *C. deuterogattii*

Like the centromeres of the closely related species *C. neoformans*, our studies show that the centromeres and pericentric regions of *C. deuterogattii* are enriched for the heterochromatic mark H3K9me2 [25]. This feature is also observed in several other fungi, including *Neurospora crassa* and *Fusarium fujikuroi*, in which centromeres are enriched for H3K9me2/3 [44,45]. Similar observations have been made in the oomycete *Phytophthora sojae* [46]. This is in contrast to *S. pombe*, in which the CENP-A-enriched regions lack heterochromatin, but pericentric regions are enriched for heterochromatin [19,38,39]. A few other fungi, such as *C. albicans* and *S. cerevisiae*, lack the H3K9me2/3 modification in their genomes. The regional centromeres of the fungal plant pathogen *Zymoseptoria tritici* lack heterochromatic marks and span actively expressed genes, despite the presence of heterochromatic marks in other regions of its genome [47].

Unlike native centromeres, neocentromeres in *C. deuterogattii* lack all heterochromatic marks and span genes (Figure S5). Other fungal neocentromeres have been studied mainly in *C. albicans* and *S. pombe*, and these neocentromeres span genes [33,34,38]. Interestingly, in both these cases, neocentromeres silence underlying genes suggesting some level of heterochromatinization in these regions. Beyond fungi, neocentromeres have been studied in a number of additional species. The native centromeres of chickens can be repetitive or non-repetitive, and heterochromatin enrichment depends upon the presence of centromeric repeats [31]. Similar to their non-repetitive native centromeres, chicken neocentromeres lack repeats and are not enriched for any of the histone marks tested [31]. Centromere repositioning occurs in plants when evolutionarily new centromeres (ENCs) are formed; however, the majority of ENCs are formed in heterochromatic regions flanking the native centromere [48]. These studies suggest that while heterochromatin might play a role in defining the location of neocentromeres in some species, it is dispensable for neocentromere formation and function in others. Together, neocentromere properties seem to be highly species-specific, similar to native centromeres which vary dramatically in their structure and properties despite a conserved function, a phenomenon known as “centromere paradox” [49].

### The heterochromatin machinery differs between closely related species *C. deuterogattii* and *C. neoformans*

5mC DNA methylation and H3K9me2 are present at centromeres in many fungal and plant species. The pericentric regions of *N. crassa* are enriched for 5mC, while H3K9me3 is found at both pericentric and CENP-A-enriched regions [44]. *S. pombe* lacks 5mC DNA methylation but H3K9me2 is present in the pericentric regions. In *C. neoformans*, both 5mC and H3K9me2 are enriched throughout centromeres, overlapping with CENP-A bound regions and also in the flanking ORF-free regions. Interestingly, we observed a different pattern between these modifications in *C. deuterogattii* where H3K9me2 was enriched through ORF-free regions but 5mC was present only in a few specific regions. Furthermore, 5mC DNA methylation was not observed on all centromeres in *C. deuterogattii* suggesting a divergence as compared to the sister species *C. neoformans*.

A recent study in *C. neoformans* revealed that deletion of *CLR4*, which encodes the enzyme that generates H3K9me2, leads to reduced 5mC levels but does not abolish them completely [26]. The authors of that study further showed that H3K9me2 regulates DNA methylation levels through two different pathways: the first pathway depends on binding of the N-terminal chromodomain of the DNA methyltransferase Dnmt5 directly to H3K9me2, and the second pathway that directs 5mC DNA methylation involves the HP1 (Heterochromatin protein 1) ortholog Swi6, which recruits Dnmt5 to H3K9me2-enriched regions. In the double mutant, in which *SWI6* is deleted and the N-terminal chromodomain of Dnmt5 is mutated, 5mC levels are similar to those in the *clr4*Δ mutant [26]. These results suggest that H3K9me2 is not the only determining factor for DNA methylation in *C. neoformans*. This hypothesis is supported by our results that H3K9me2 and 5mC can be enriched independent of one another in some regions of the genome and localize with each other in other regions. Interestingly, Swi6 is present in *C. deuterogattii*, and future studies could focus on the interaction of these proteins in this species and how this may differ from their interactions in *C. neoformans*. Moreover, the exact roles of 5mC and H3K9me2 modifications are not yet understood in either of these two species and it is possible that they play different roles in different species.

### Neocentromere formation in *C. deuterogattii* is defined by “CENP-A cloud”

In *S. pombe*, neocentromeres form near heterochromatic repeats in sub-telomeric regions [38], whereas in *C. albicans* and chicken DT40 cells, neocentromeres most often form in close proximity to native centromeres, likely due to the seeding of CENP-A near the native centromere i.e. the “CENP-A cloud” [32–34,36]. The “CENP-A cloud” not only influences the chromosomal regions directly flanking the centromere, but also chromosomal regions that are in close vicinity to the centromere due to chromosome folding, and these regions can be several kilobases away from the native centromeric chromosomal location [50]. Previously, we showed that most neocentromeres of *C. deuterogattii* chromosomes 9 and 10 form in close proximity to the deleted native centromere [40]. The neocentromere formation of chromosome 4 in this study shows that neocentromeres are formed in close proximity to the native *CEN4* suggesting a possible role of “CENP-A cloud” in this species as well.

This hypothesis was further supported by the observation that deletion of native centromere in a fusion chromosome, that harbored neocentromere previously, also results in the formation of a novel neocentromere next to the native centromere. These results also suggested that neocentromeres lacks an epigenetic memory and the location of native centromere influences neocentromere formation in this species. However, this hypothesis needs further testing by using a more robust system as well as with neocentromeres in other chromosomes in *C. deuterogattii* and should be the subject of further investigation.

### Dicentric chromosomes are unstable in *C. deuterogattii*

Naturally occurring dicentric B-chromosomes in maize (*Zea mays*) are stabilized by the inactivation of one of the centromeres [51]. Interestingly the inactive centromere can be reactivated due to chromosome recombination, which suggests there is an epigenetic memory or at least a preference for the centromere location and chromosomal context. Dicentric chromosomes in *S. pombe* have been obtained by chromosomal fusion of chromosomes 2 and 3 [52]. This study revealed multiple fates for these dicentric chromosomes: 1) epigenetic inactivation of a centromere, resulting in a stable, fused chromosome; 2) stabilization of the chromosome fusion by deletion of a centromeric region or; 3) chromosome breakage resulting in three chromosomes with functional centromeres. Furthermore, the inactivation of a centromere was found to be associated with the spreading of heterochromatin into the CENP-A-bound central core.

All of these examples describe chromosome fusions resulting in stable chromosomes with only one active centromere. Stable dicentric chromosomes in human cell lines have been engineered by displacement of the telomere protein Trf2 [53]. Half of the isolates obtained were *bona fide* dicentrics and stable for >150 cell divisions. In the other half of the isolates, the chromosome was stabilized either by partial deletion of centromeric repeats in one centromere or via epigenetic inactivation. In our study, we obtained two colonies that underwent chromosome-chromosome fusion induced via CRISPR-CAS9. Analysis using PFGE and nanopore sequencing suggested that one of two strains had a stable chromosome fusion, whereas the fused chromosome in the second isolate broke before the dicentric nature could be analyzed. Analysis of the stable fused chromosome using ChIP-qPCR indicated that only one or the other centromere was active and the chromosome was most likely not a true dicentric. This result was similar to the result in our previously described fused chromosome, which formed due to the inactivation of the neocentromere of chromosome 10 [40]. In that case, the neocentromere was inactivated after the chromosome fusion and only the active native centromere of the partner chromosome remained functional. Overall, these results suggest that dicentric chromosomes in *C. deuterogattii* are not stable independent of whether they originate from neocentromeres or artificial chromosome-chromosome fusion.

### The centromeric *URA5* transgene lacks heterochromatin silencing

Some species have native centromeres that span actively expressed genes, and human artificial chromosomes have centromeres that are only sparsely enriched for heterochromatin [47,54]. Neocentromeres in *C. deuterogattii* and DT40 cells can also span actively expressed genes, whereas neocentromeres in *C. albicans* and *S. pombe* silence the underlying genes [32,34,38,40]. Along similar lines, when a transgene is inserted into the native centromeres of *C. albicans* or *S. pombe*, the gene is silenced. In *S. pombe*, the extent of transgene silencing depends on the location of the insertion, and the transgene is not silenced when integrated into a CENP-A-binding region [55,56]. On the other hand, when the *URA3* gene is inserted into a centromere of *C. albicans*, the gene is expressed conditionally and exhibits reversible silencing [37]. Moreover, the transgene in *C. albicans* is bound by CENP-A when it is silenced but not when it is expressed. In a previous study, we found that neocentromeric genes are expressed in *C. deuterogattii* [40]. In contrast to that, here we showed that integration of a transgene into a native centromere in *C. deuterogattii* also led to a shift of CENP-A binding. Specifically, the *URA5* gene integrated into *CEN2* is expressed at the same levels as that of *URA5* in the native locus. Furthermore, the *URA5* gene inserted into *CEN2* was not decorated with H3K9me2. Our ChIP-seq experiments for CENP-A and H3K9me2 conducted to characterize this phenomenon were performed with cells grown without selection for *URA5* expression, providing evidence for the stability of the transgene and its expression status. However, we cannot exclude the possibility that CENP-A binding might revert back to the native *CEN2* position and that H3K9me2 would span the *URA5* gene if the strain were grown for many generations in media without selection.

### Centromeric drift in *C. deuterogattii*

Several studies have shown that CENP-A peaks drift (or shift) within a defined chromosomal region. For example, CENP-A profiles of several maize isolates showed that centromeric drift occurs in centromeres 5 and 8 [57]. Centromeres 4 and 7 in donkeys have two distinct CENP-A peaks; however, hybrid mules derived from donkeys and horses have only one CENP-A-enriched region in both centromeres [57]. Compared to donkey centromeres 4 and 7, the CENP-A-enriched regions of mule centromeres are slightly shifted and not completely homologous to the parental donkey CENP-A locations. ChIP-seq of five independent horses revealed five unique CENP-A-bound regions for the non-repetitive centromere 11 [58]. Additionally, the CENP-A-enriched region of the non-repetitive centromere Z in chicken DT40 cells drifts in the population, and this drift occurs more frequently in CENP-U and CENP-S mutants [43].

We observed a similar phenomenon of centromere drift in *C. deuterogattii*. Specifically, we identified drifting of CENP-A binding in two different centromeres: *CEN5* and *CEN14*. Centromere drift has not been observed in any fungal species yet, and this is possibly due to a lack of multiple ChIP-seq datasets for a single species. Interestingly, we observed that drift in *CEN5* was associated with the presence of fragments of a specific transposon, which is present only in a few copies throughout centromeres. No such defining factor has been identified for centromere drift in other organisms suggesting more large-scale ChIP-seq studies will be required to identify such determining factor.

## Material and methods

### Strains, primers, and culture conditions

Primers are listed in Supplementary Table 1. Strains used in this study are listed in Supplementary Table 2. All strains were stored in glycerol stocks at -80°C, inoculated on solid YPD (yeast extract, peptone, and dextrose) medium, and grown for two days at 30°C. Liquid YPD cultures were inoculated with single colonies from solid medium and grown, while shaking, at 30°C overnight.

### Genetic manipulations

Prior to biolistic transformation or CRISPR-Cas9-mediated transformation, PCR products with homologous DNA sequences (1 to 1.5 kb) flanking the deleted region were PCR-amplified with Phusion High-Fidelity DNA Polymerase (NEB, Ipswich MA, USA). These flanking PCR products were fused to two sides of either a *NEO* or *NAT* dominant selectable marker via overlap PCR, conferring G418 or nourseothricin resistance, respectively. For the deletion of *CEN4*, two guide RNAs (gRNAs) flanking *CEN4* were designed. The chromosome fusion of chromosome 4 and 10 was also mediated by two gRNAs targeting the sub-telomeric region of chromosome 4 and the sub-telomeric region of chromosome 10, respectively. Biolistic transformation and CRISPR-Cas9-mediated transformation were performed as previously described, and transformants were selected on YPD medium containing G418 (200 μg/mL) or nourseothricin (100 μg/mL) [40,59,60]. All mutations were introduced into a previously generated *C. deuterogattii* strain that expresses mCherry-CENP-A. To confirm the correct replacement of the region of interest by the appropriate drug resistance marker, PCR analysis was conducted for the 5’ junction, 3’ junction, and spanning the target locus.

To delete the *DMT5* gene, 1-kb homologous flanking sequences were utilized to make a deletion allele with a *NEO* selectable marker in a similar approach as described above. Two independent gRNAs cleaving the *DMT5* gene at two different locations were used along with the recombination template for the CRISPR-Cas9-mediated transformation. Transformants obtained from two independent biolistics transformation experiments were screened and confirmed with 5’ and 3’ junction PCRs, resulting in two independent *dmt5*Δ mutants from each biolitstics transformation.

Prior to the integration of *URA5* in centromere 2 (*CEN2*), an mCherry-CENP-A tagged wild-type strain was plated on a medium containing 5-FOA, and spontaneously-derived resistant colonies were isolated. These colonies were further screened for their lack of growth on SD-uracil drop-out medium and normal growth on 5-FOA-containing medium. Colonies growing on 5-FOA but not on SD-uracil were identified as *ura5* mutant colonies and utilized for further experiments. One such 5-FOA resistant, uracil auxotrophic isolate (KS174) was shown to be auxotrophic for uracil and sequence analysis of the *URA5* gene revealed a single base-pair deletion mutation resulting in the lack of adenine at position 156 resulting in a frameshift. This *ura5* mutant then served as the background strain in which the *URA5* gene was integrated into *CEN2*, and transformants were selected on SD-uracil medium. The integration of *URA5* at the correct location was then confirmed by PCR analysis and Southern blot hybridization (as shown in Figure S9).

For the dicentric isolates, chromosome fusion was mediated by CRISPR-Cas9 with two guide RNAs targeting the sub-telomeric regions of chromosome 4 and 10, and the resulting double-stranded breaks were repaired using a linear overlap PCR product that contained a 1.5 kb region homologous to chromosome 4, a *NAT* selection marker, and a 1.5 kb region homologous to chromosome 10.

### Chromatin immunoprecipitation (ChIP) followed by high-throughput sequencing or qPCR

ChIP analyses were performed as previously described [40]. A polyclonal antibody against mCherry (ab183628, Abcam) was used to identify the CENP-A-enriched regions. To identify histone H3K9me2 and H3K4me2-enriched regions, ChIP analysis was conducted with a monoclonal antibody against histone H3K9me2 (ab1220, Abcam) or H3K4me2 (#39141, Active Motif). Subsequently, the samples were analyzed by qPCR or subjected to the library preparation and Illumina sequencing at the Duke University Sequencing and Genomic Technologies Shared Resource facility. Sequencing was performed with a NovaSeq 6000 sequencer and 50/100-bp PE reads were obtained. qPCRs were performed in triplicate with Brilliant III Ultra-Fast SYBR Green qPCR Master Mix (Agilent Technologies) on an ABI 1900HT qPCR machine.

### Data analysis

Illumina reads obtained after ChIP-seq generated for this study, and previously published ChIP-seq datasets, were processed and analyzed following the protocol described below [21,40]. Illumina reads were filtered and trimmed as previously described [61]. ChIP-seq sequencing was mapped to the *C. deuterogattii* R265 reference genome by bowtie2 with standard settings [62]. Subsequently, the reads were processed by using Samtools and Bedtools [63,64]. BamCompare with standard settings was used to normalize the samples with the input and, subsequently, the data was visualized with the Integrative Genome Viewer (IGV) and IGV tools [64,65]. Macs2 was used for statistical tests with standard settings [66]. Hisat2 was used to remap a previously generated RNA-seq dataset (NCBI Sequence Read Archive, SRR5209627) to the *C. deuterogattii* R265 reference genome [67][68]. Command-line scripts used to process, map and analyze the data are shown in supplementary file S1.

### Verification of *URA5* gene activity in wild-type and *URA5*in*CEN2* mutant

Wild-type and *URA5*in*CEN2* mutant from -80°C stocks were inoculated on solid YPD and incubated at 30°C. Ten single colonies of each isolate were isolated from the YPD plate, inoculated in liquid YPD cultures and grown overnight at 30°C. Overnight cultures were washed twice with water and serial dilution series were performed. Cells were inoculated on solid YPD with a dilution of 10^5^ and a dilution of 10^2^ was used to inoculated YNB plates containing 5FOA. Plates were incubated at 30°C until colonies appeared, subsequently all colonies were counted and the mutation rate was calculated as previously described [69].

### Pulsed-field gel electrophoresis (PFGE)

Isolation of whole chromosomes and conditions for PFGE analysis was performed as previously described [70].

### Nanopore sequencing and 5mC analysis

The DNA preparation for nanopore sequencing was done using CTAB based method as described previously [71]. The DNA obtained was checked for quality and size using NanoDrop and PFGE, respectively. Using the high-quality DNA, sequencing was performed on the MinION device where all the strains sequenced were multiplexed together in a single flow cell. The samples were barcoded using the EXP-NBD103 kit and libraries were made using the SQK-LSK109 kit as per the manufacturer’s instructions. The libraries generated were sequenced on the R9.4.1 flow cell, reads were obtained in FAST5 format, which was then converted to .fastq using Guppy_basecaller (v 4.2.2_linux64). The Fastq reads were de-multiplexed using Guppy_barcoder (Part of Guppy_basecaller) and barcodes were trimmed during this processing. The reads, thus obtained for each sample were then used for calling methylation against the reference R265 genome suing nanopolish (v 0.13.2) [72]. The methylation frequency was calculated using the calculate_methylation_frequency.py script available with nanopolish and then converted to bedGraph for visualization purposes in IGV.

### Bisulfite sequencing

Genomic DNA was isolated following the CTAB method, checked for quality, and submitted to the Duke University Sequencing and Genomic Technologies Shared Resource facility for whole-genome bisulfite sequencing [73]. After sequencing, the reads were analyzed and mapped to the *C. deuterogattii* R265 reference genome using Bismark (https://www.bioinformatics.babraham.ac.uk/projects/bismark/). The default parameters for alignment and methylation extraction were used for both the wild-type strain and the *dmt5*Δ mutant. Methylation extraction results were obtained as a .bed file that was then imported into IGV for visualization.

### Deposited data

ChIP sequences have been deposited under NCBI BioProject Accession ID: ####.

## Acknowledgements

We thank Shelby Priest for her comments on the manuscript. These studies were supported by NIH/NIAID grants R01 AI050113-17 and R37 MERIT award AI039115-24 to JH. JH is co-director and fellow of the CIFAR program the Fungal Kingdom: Threats & Opportunities.

## Supplementary figures

**Figure S1.**
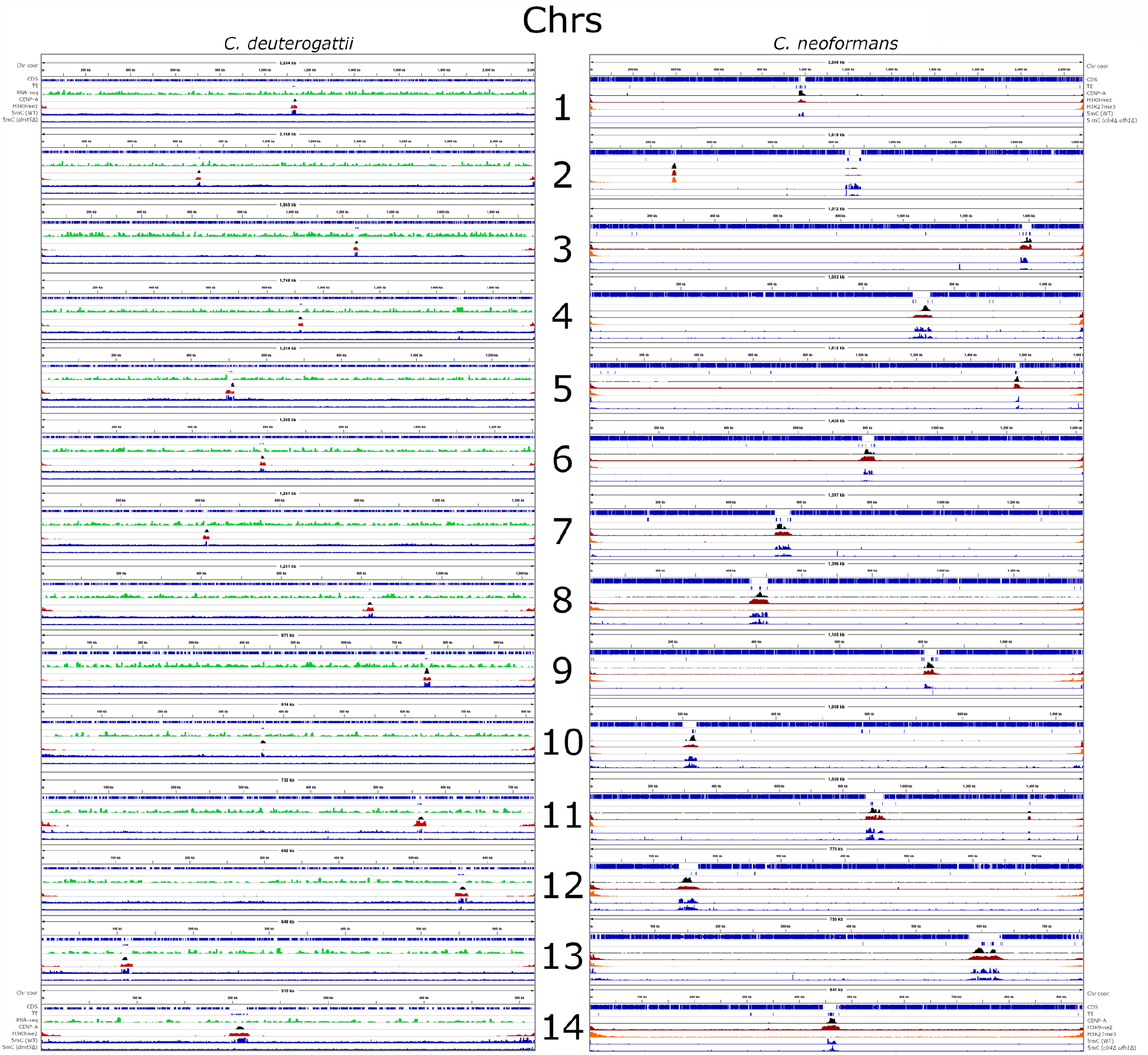
Whole-genome view of histone marks enrichment and 5mC DNA methylation. All 14 chromosomes are depicted for both *C. neoformans* and *C. deuterogattii*. For each chromosome, plots presented show the chromosome coordinates, gene content/CDS (blue), TE content (blue), RNA-seq (for *C. deuterogattii* only) (green), CENP-A enrichment (black), H3K9me2 enrichment (red), H3K27me3 enrichment (for *C. neoformans* only) (orange), and 5mC data (blue) [25].

**Figure S2.**
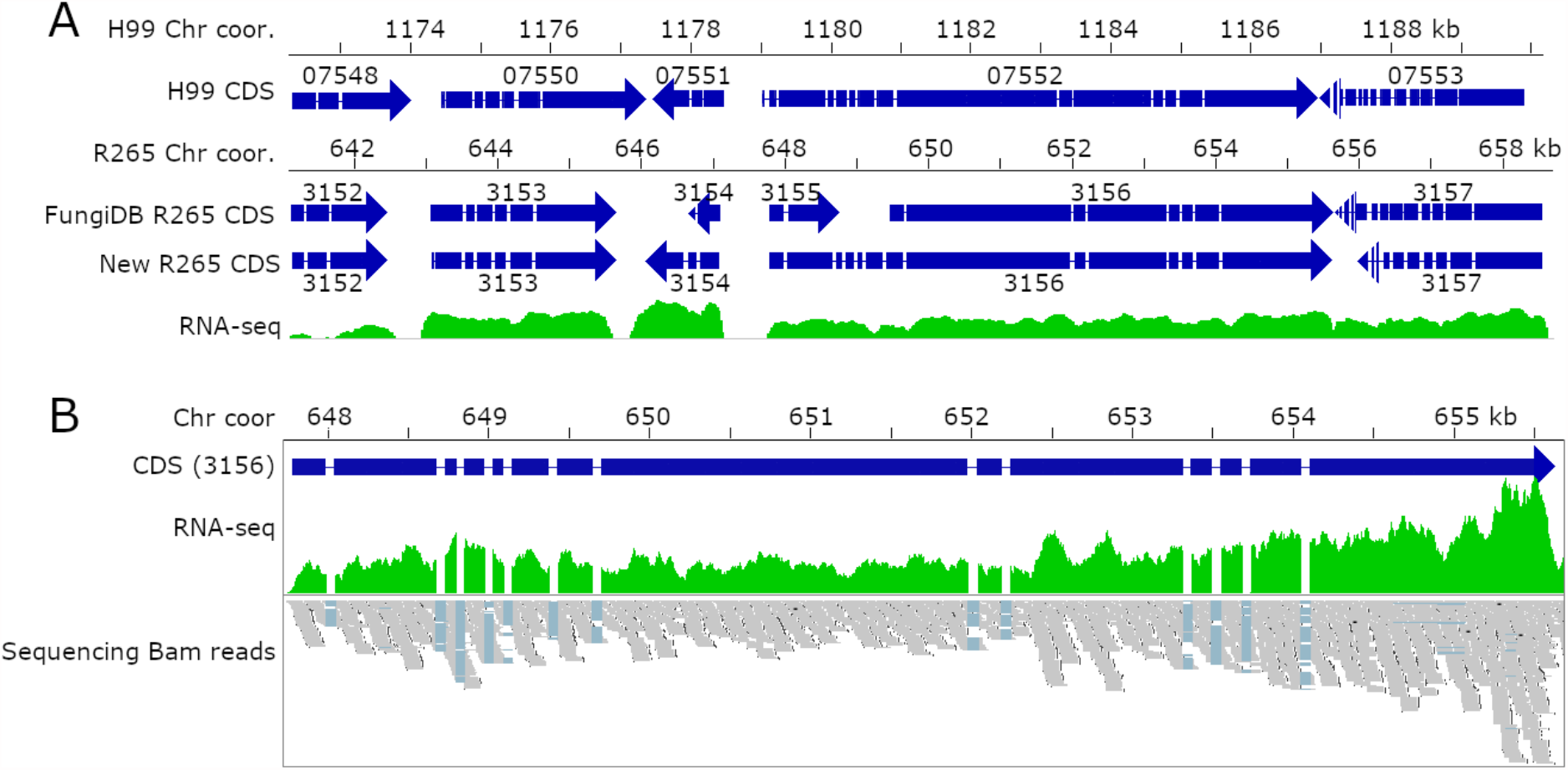
The *C. deuterogattii DMT5* gene encoding Dnmt5 is intact and expressed similarly to the *C. neoformans* ortholog. **(A)** Alignment view with genomic regions surrounding the *DMT5* gene (CNAG_07752 and CNBG_3156) of *C. neoformans* and *C. deuterogattii*. Shown at the top of the panel are the chromosomal coordinates of *C. neoformans*. The genes are indicated with blue arrows with exons and introns marked. For *C. deuterogattii*, two gene annotations are shown. The old (FungiDB) annotation predicted that *DMT5* was truncated. The new genome annotation shows that the *DMT5* gene is full-length and has a similar length to the *C. neoformans* ortholog [41]. **(B)** Detailed view of *C. deuterogattii* introns and exons of the *DMT5* gene is supported by RNA-seq reads, which are shown to support the gene structure.

**Figure S3.**
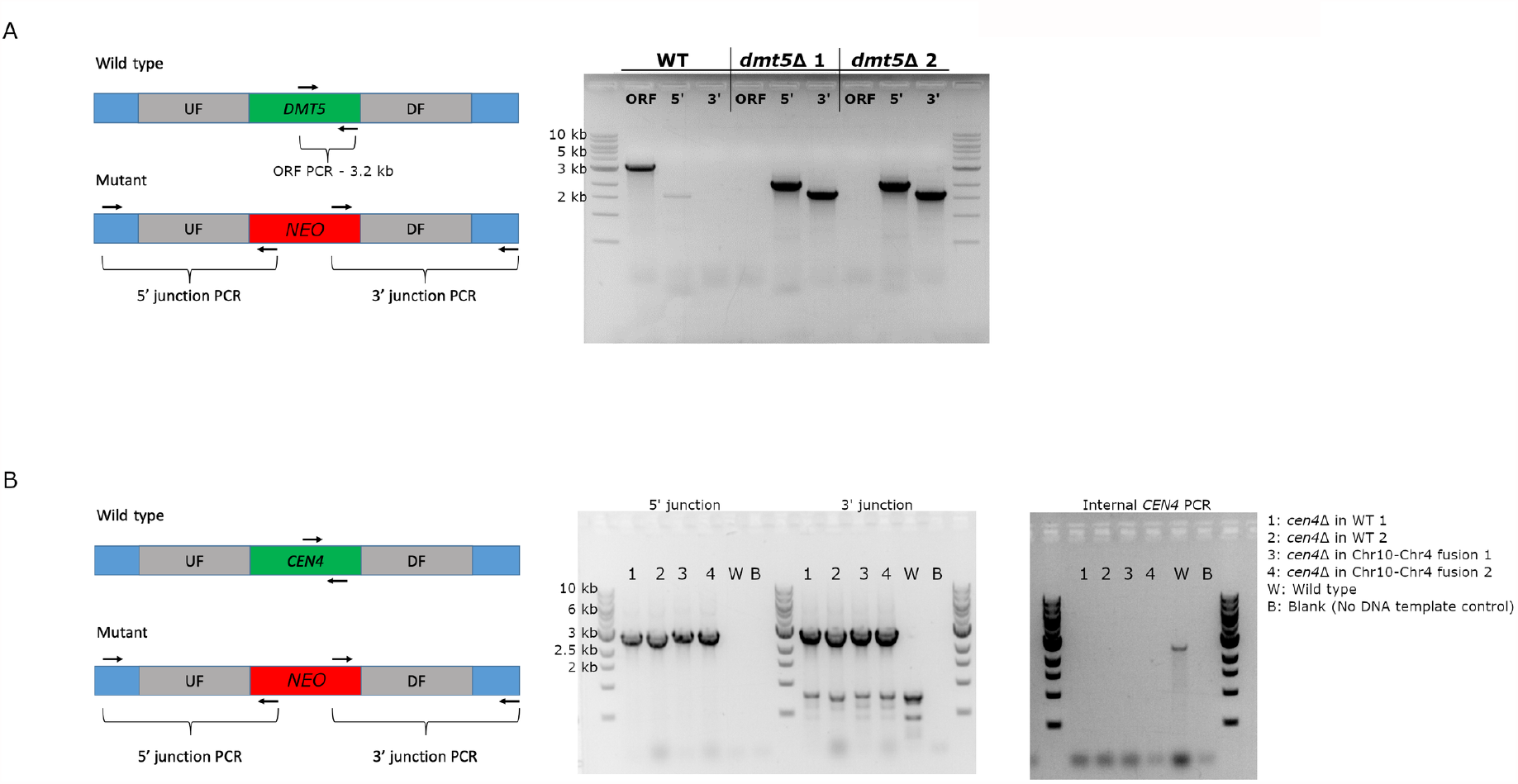
PCR confirmation of gene deletion mutants. PCR confirmations for *dmt5*Δ and *cen4*Δ mutants are shown. Panel **(A)** shows PCR confirmations for *dmt5*Δ; panel **(B)** shows PCR confirmations for *cen4*Δ. For both mutants, the 5’ and 3’ junction, as well as internal/ORF PCRs, are shown.

**Figure S4.**
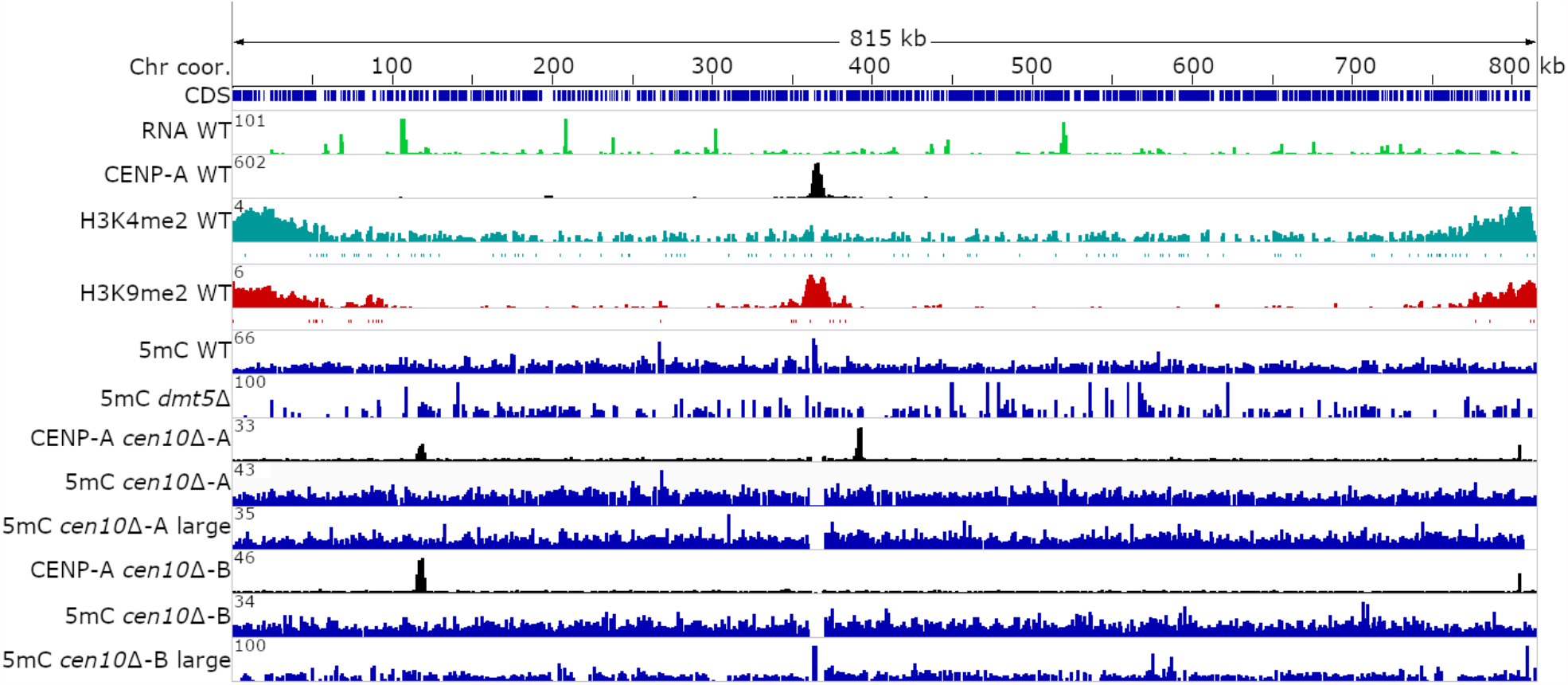
Neocentromeres are not enriched for 5mC DNA methylation. ONT sequencing was performed for two previously obtained *cen10*Δ and the derived chromosome fusions (large) that have silenced neocentromeres. The complete chromosome 10 is shown and chromosomal coordinates are indicated. The wild-type genome annotation (CDS), RNA-seq (green), CENP-A ChIP-seq (black), H3K4me2 ChIP-seq (blue) and ChIP-seq for H3K9me2 (red) is shown. DNA methylation analyses based on ONT sequencing are shown in blue for WT, *dmt5*Δ, and *cen10*Δ mutants -A and -B and their derived chromosome fusion products. For each neocentromere mutant, the CENP-A ChIP-seq track shows the neocentromere location.

**Figure S5.**
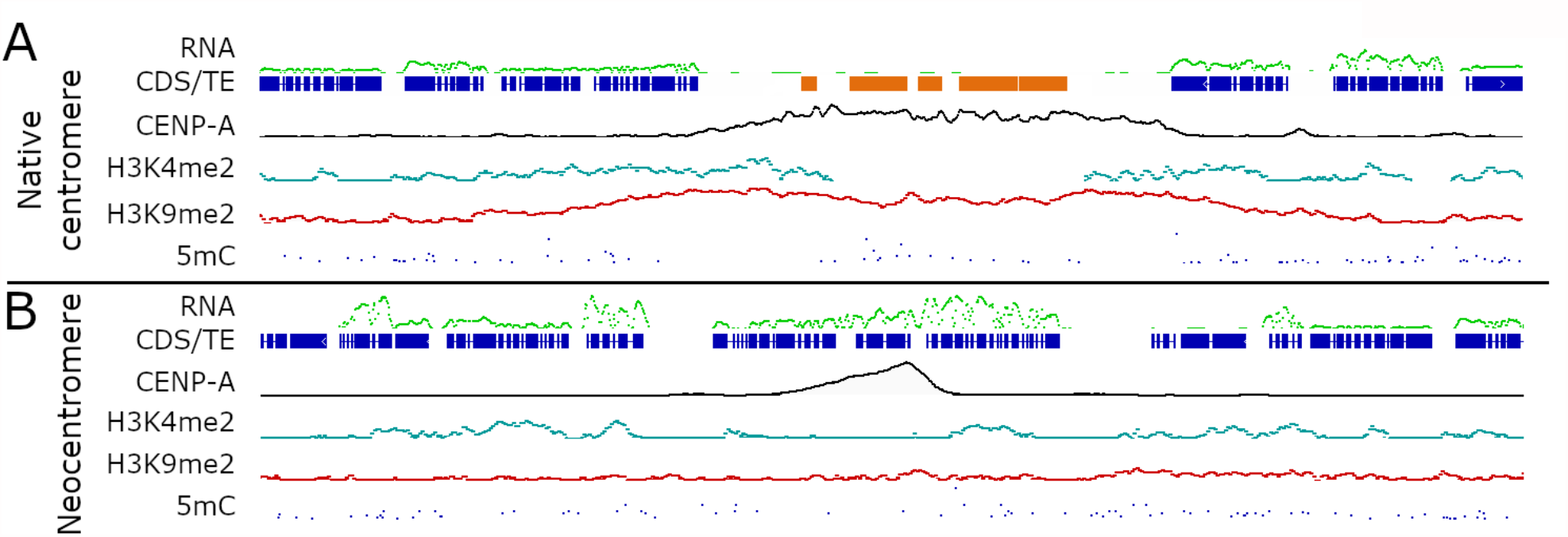
Schematic overview of *C. deuterogattii* native and neocentromeres. Panel **(A)** shows the organization of a native centromere. Native centromeres are located in ∼14.5 kb ORF-free chromosomal regions and these regions are enriched for truncated transposable elements (TE, orange). The whole centromeric region is modestly enriched for 5mC DNA methylation (blue), enriched for H3K9me2 (red) and within this heterochromatic region, the CENP-A peak (Black) is located. The centromeric regions are flanked by genes (dark blue) that are actively expressed (green) and enriched for the euchromatic histone mark H3K4me2 (green). Panel **(B)** shows an example of a neocentromere. Neocentromeres span actively expressed (green) genes (blue), are determined by the presence of CENP-A (black) and lacked any enrichment for the epigenetic marks that were analyzed, while the genes are still enriched for H3K4me2 (green).

**Figure S6.**
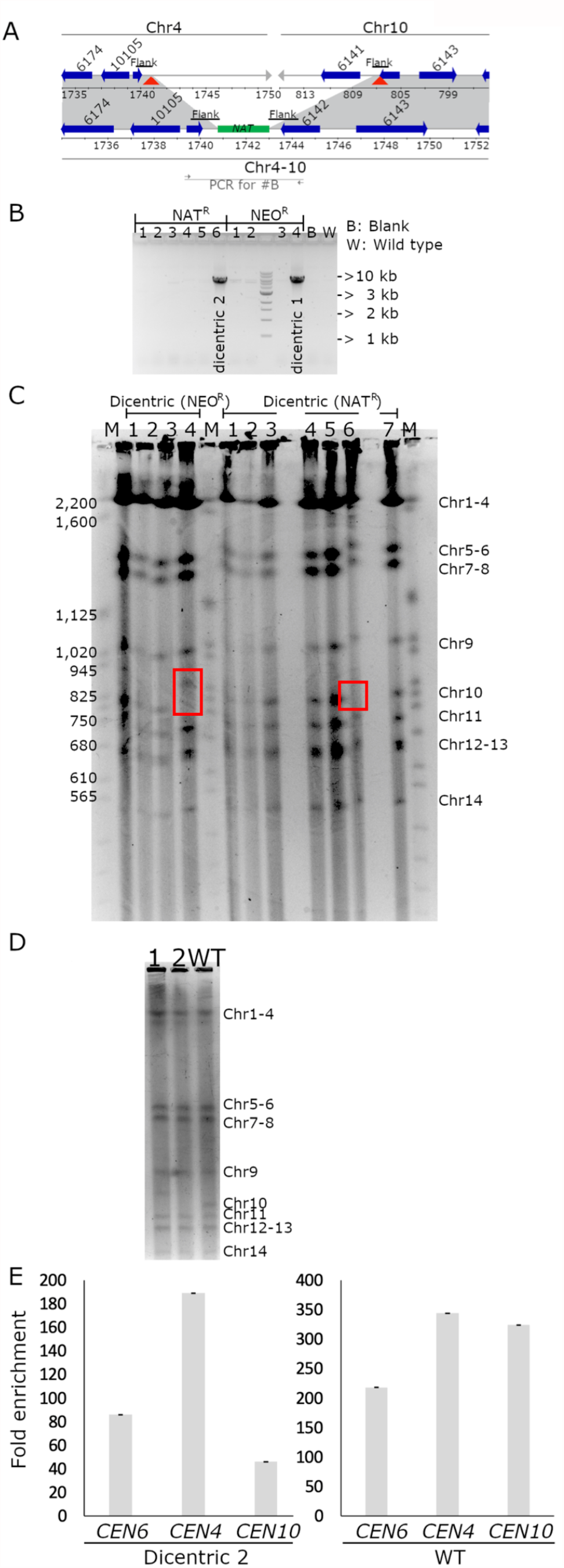
Formation of a dicentric chromosome. **(A)** Schematic view of the formation of a dicentric chromosome. At the top, a sub-telomeric region of wild-type Chr4 and Chr10 is shown. The chromosomal targets of the guide RNAs are depicted by red triangles. The double-stranded DNA breaks were repaired by homologous recombination that was mediated by an overlap PCR product containing regions homologous to both chromosomes flanking a selectable marker. The homologous regions are depicted by black lines and labeled “Flank”. **(B)** Spanning PCR analysis confirmed chromosome fusion in mutant 1 (G418^R^ 4) and mutant 2 (Nat^R^ 6) are shown. The region amplified for the spanning PCR is indicated in panel A with a bar labeled with “PCR for #B”. **(C)** PFGE analysis with all mutants obtained after recovering mutants on selective media. Dicentric 1 (G418^R^ 4) and 2 (Nat^R^ 6) lack a wild-type size band for Chr10 confirming the spanning PCR products in panel A and these are indicated with a red rectangle. **(D)** PFGE analysis shows that wild-type Chr 10 is absent in dicentric strains 1 and 2, which confirms that Chr4 and Chr10 are fused in these two strains. Instead of the wild-type Chr10 band, mutant dicentric 1 has an additional band (∼60 kb higher) than the wild-type Chr10 band. Based on this PFGE, dicentric 1 has no additional resolvable bands. **(E)** ChIP-qPCR analyses were performed for dicentric mutant 2 and wild type. CENP-A fold enrichment is shown for *CEN4* and *CEN10* and a positive control (*CEN6*). The fold-enrichment was compared to actin as the negative control.

**Figure S7.**
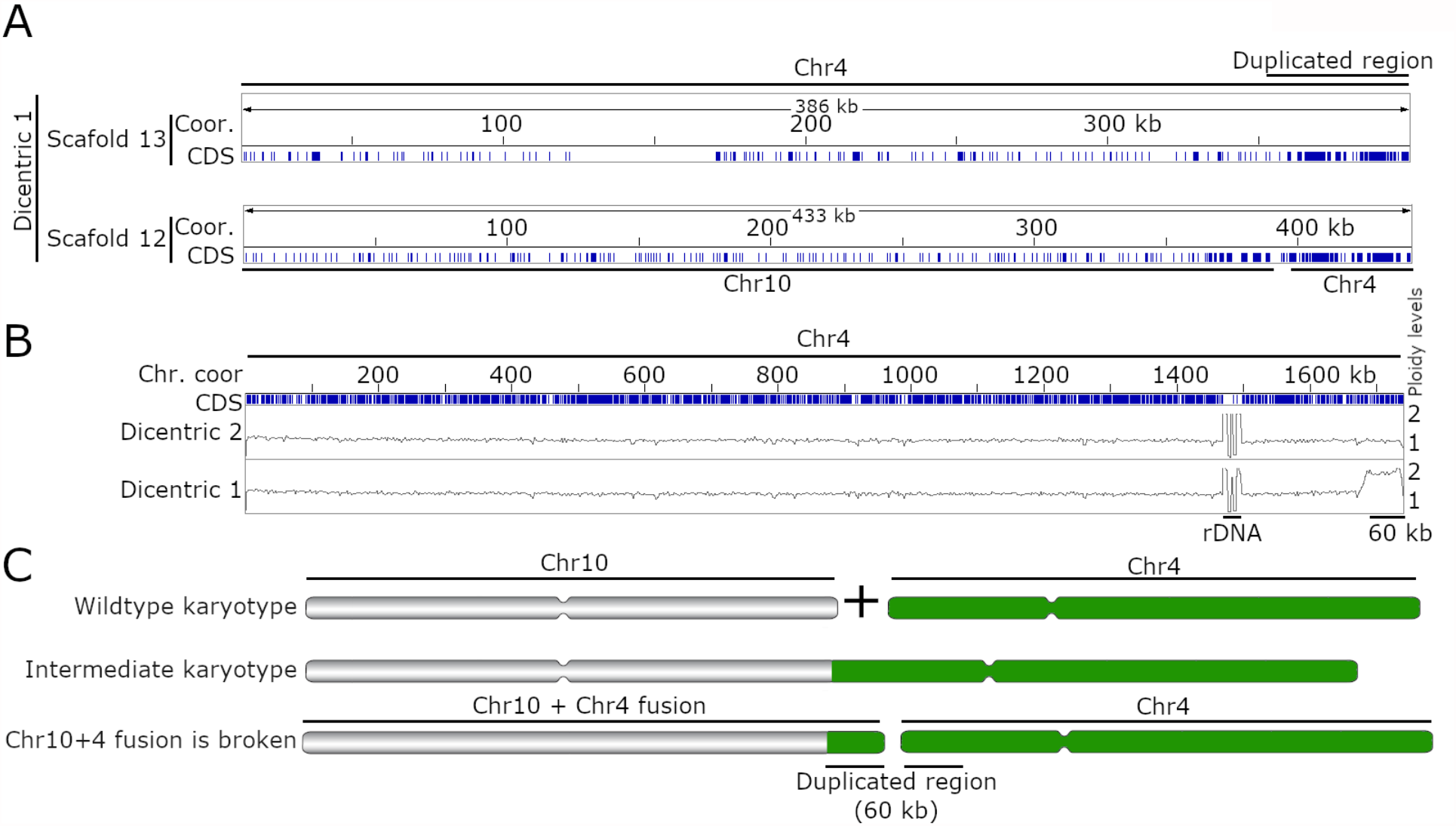
Nanopore sequencing reveals that the fused dicentric chromosome underwent chromosomal breakage. Panel (**A**) shows the two scaffolds that harbor the ∼60 kb duplicated region. Scaffold 13 corresponds to a part of chromosome 4, and scaffold 12 is the broken chromosomal fusion product of chromosome 10 and 4. (**B**) The panel shows Illumina sequencing of the dicentric mutants mapped to the wild-type genome. Short read sequencing indicated that a ∼60 kb region of chromosome 4 is duplicated in dicentric 2 and has a ploidy level of two as compared to the rest of the chromosome. As a control, dicentric isolate 2 was sequenced and this strain has a ploidy level of one for the entire chromosome 4. (**C**) Putative model explaining the chromosome fusion between chromosome 4 and 10, the intermediate state in which the fused chromosome is intact, and the final karyotype after the chromosomal breakage for dicentric 1.

**Figure S8.**
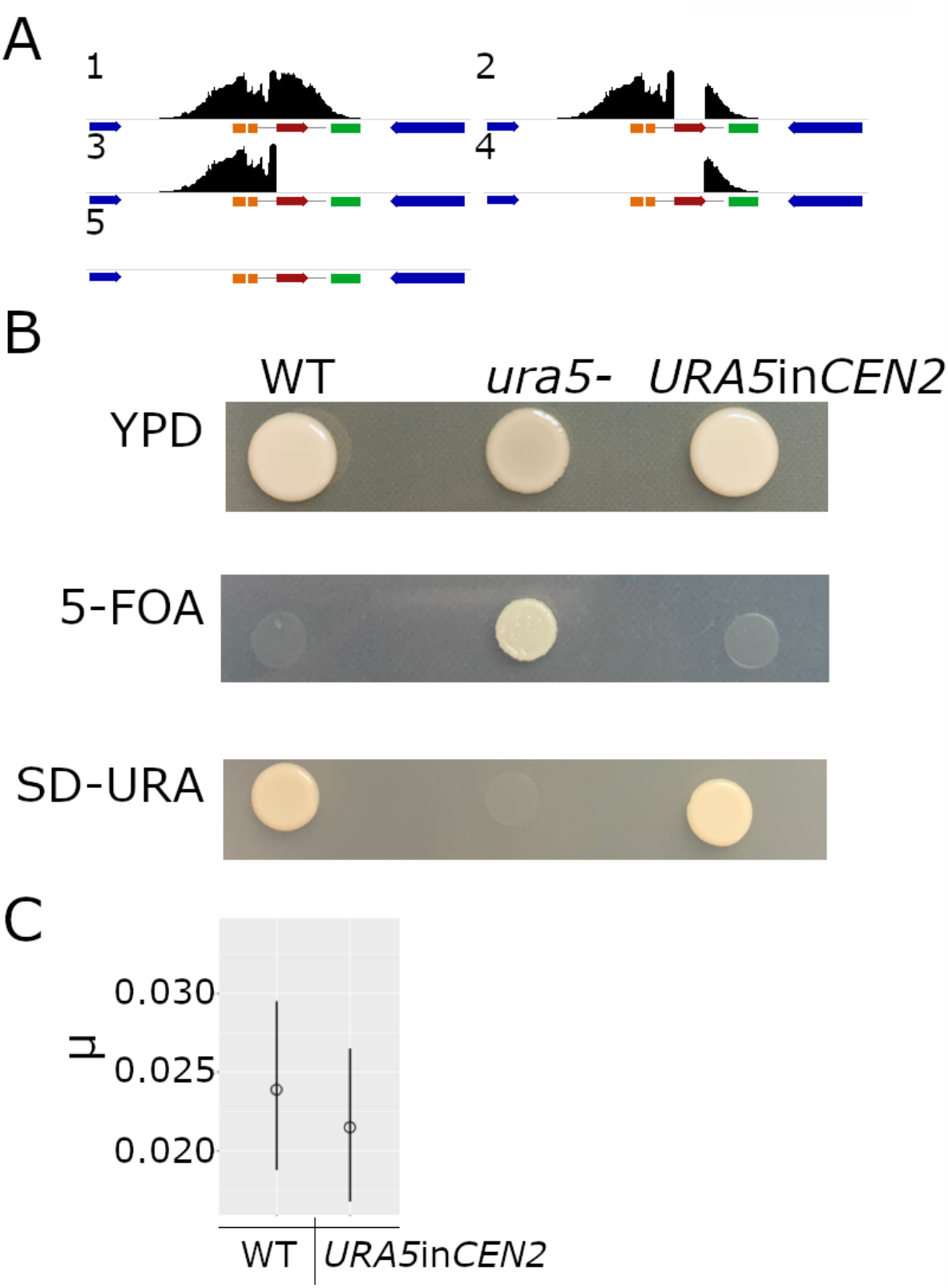
*URA5* integration into *CEN2*. **(A)** A schematic model of *CEN2* is shown. The CENP-A peak is shown in black, genes flanking the pericentric region are shown in blue. The *URA5* gene is shown in red and the truncated transposable elements are shown in orange and green (Tcn4 and Tcn6). Hypothetical outcomes for *URA5* integration into *CEN2* include: (1) CENP-A would cover the *URA5* gene making the CENP-A-bound region larger than the native centromere; (2) *URA5* gene might divide the CENP-A-enriched region into two independent regions; (3 & 4) either one of the regions flanking *URA5* would be enriched for CENP-A, generating a smaller centromere; (5) *URA5* integration abolishes *CEN*2 function leading to neocentromere formation. **(B)** Prior to the ChIP-seq experiment for *URA5*in*CEN2*, the strain was tested for *URA5* expression. For all three plates, the wild type was included as a control, and *ura5-*is the parental strain in which the *URA5* gene was integrated into *CEN2*. As expected all three strains grew on the control medium (YPD). Only *ura5-*strain was able to grow on a medium containing 5-FOA. On the SD-uracil (SD-URA) medium, the wild-type and *URA5*in*CEN2* strains were able to grow due to the presence of an active *URA5* gene. **(C)** Assay shows that the *URA5* gene in wild-type and *URA5*in*CEN2* mutant has a similar mutation rate when growing on YPD or 5-FOA medium. The mutation rate (μ) is shown on the y-axis.

**Figure S9.**
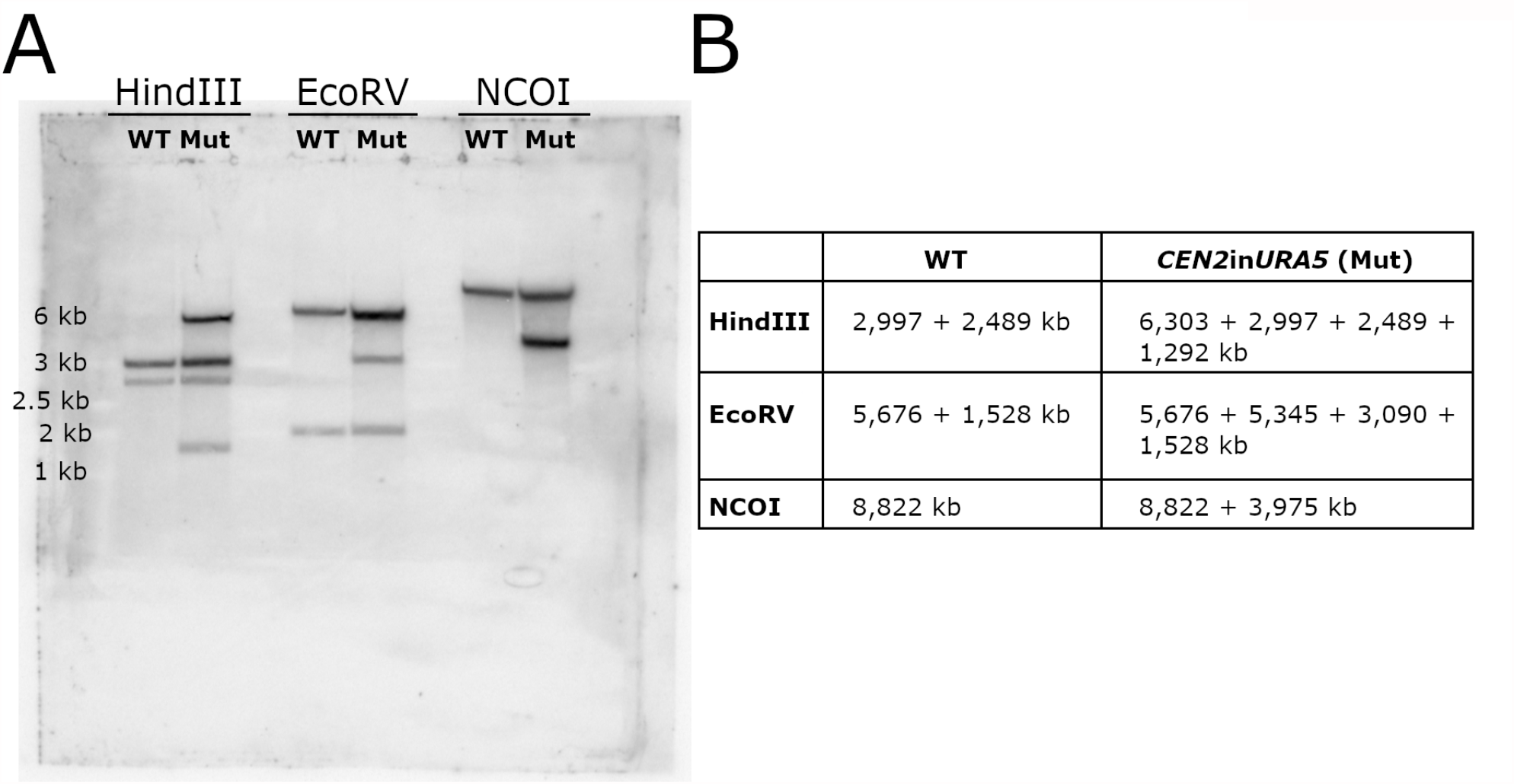
Southern blot analysis confirms *URA5* integration into *CEN2* in the genome. **(A)** Southern blot analysis with three independent restriction digest analyses of the *CEN2*in*URA5* strain. The *URA5* gene sequence was used as a probe. The Southern blot hybridization pattern confirms that a single copy of the *URA5* gene has been inserted at the desired targeted site in *CEN2*. (**B**) Table indicating the expected restriction digest product sizes. Because the *CEN2*in*URA5* strain still has the *ura5*-gene present at its native locus, the pattern shows products from both the *ura5* native gene and the *CEN2*in*URA5* transgene.

## References

1. Barra V, Fachinetti D. The dark side of centromeres: types, causes and consequences of structural abnormalities implicating centromeric DNA. Nat Commun. 2018;9: 4340. doi:10.1038/s41467-018-06545-y

2. Freitag M. The kinetochore interaction network (KIN) of ascomycetes. Mycologia. 2016;108: 485–505. doi:10.3852/15-182

3. Navarro-Mendoza MI, Pérez-Arques C, Panchal S, Nicolás FE, Mondo SJ, Ganguly P, et al. Early diverging fungus Mucor circinelloides lacks centromeric histone CENP-A and displays a mosaic of point and regional rentromeres. Curr Biol. 2019;29: 3791-3802.e6. doi:10.1016/j.cub.2019.09.024

4. Drinnenberg IA, DeYoung D, Henikoff S, Malik HS. Recurrent loss of CenH3 is associated with independent transitions to holocentricity in insects. eLife. 2014;3: 1–19. doi:10.7554/eLife.03676

5. Hooff JJ, Tromer E, Wijk LM, Snel B, Kops GJ. Evolutionary dynamics of the kinetochore network in eukaryotes as revealed by comparative genomics. EMBO Rep. 2017;18: 1559– 1571. doi:10.15252/embr.201744102

6. Saldivia M, Fang E, Ma X, Myburgh E, Carnielli JBT, Bower-Lepts C, et al. Targeting the trypanosome kinetochore with CLK1 protein kinase inhibitors. Nat Microbiol. Springer US; 2020;5: 1207–1216. doi:10.1038/s41564-020-0745-6

7. Friedman S, Freitag M. Centrochromatin of fungi. Centromeres and Kinetochores. Cham, Switzerland: Springer International Publishing; 2017. pp. 85–109. doi:10.1016/S0092-8674(03)00115-6

8. Kobayashi N, Suzuki Y, Schoenfeld LW, Müller CA, Nieduszynski C, Wolfe KH, et al. Discovery of an unconventional centromere in budding yeast redefines evolution of point centromeres. Curr Biol. 2015;25: 2026–2033. doi:10.1016/j.cub.2015.06.023

9. Guin K, Sreekumar L, Sanyal K. Implications of the evolutionary trajectory of centromeres in the fungal kingdom. Annu Rev Microbiol. 2020;74: 835–853. doi:10.1146/annurev-micro-011720-122512

10. Padmanabhan S, Thakur J, Siddharthan R, Sanyal K. Rapid evolution of Cse4p-rich centromeric DNA sequences in closely related pathogenic yeasts, Candida albicans and Candida dubliniensis. Proc Natl Acad Sci U S A. 2008;105: 19797–19802. doi:10.1073/pnas.0809770105

11. Chatterjee G, Sankaranarayanan SR, Guin K, Thattikota Y, Padmanabhan S, Siddharthan R, et al. Repeat-associated fission yeast-like regional centromeres in the ascomycetous budding yeast Candida tropicalis. PLOS Genet. 2016;12: e1005839. doi:10.1371/journal.pgen.1005839

12. Mishra PK, Baum M, Carbon J. Centromere size and position in Candida albicans are evolutionarily conserved independent of DNA sequence heterogeneity. Mol Genet Genomics. 2007;278: 455–65. doi:10.1007/s00438-007-0263-8

13. Sanyal K, Baum M, Carbon J. Centromeric DNA sequences in the pathogenic yeast Candida albicans are all different and unique. Proc Natl Acad Sci U S A. 2004;101: 11374– 9. doi:10.1073/pnas.0404318101

14. Freire-Benéitez V, Price RJ, Buscaino A. The chromatin of Candida albicans pericentromeres bears features of both euchromatin and heterochromatin. Front Microbiol. 2016;7: 1–11. doi:10.3389/fmicb.2016.00759

15. Price RJ, Weindling E, Berman J, Buscaino A. Chromatin profiling of the repetitive and nonrepetitive genomes of the human fungal pathogen Candida albicans. mBio. 2019;10: 1–17. doi:10.1128/mBio.01376-19

16. Freire-Benéitez V, Price RJ, Tarrant D, Berman J, Buscaino A. Candida albicans repetitive elements display epigenetic diversity and plasticity. Sci Rep. 2016;6: 1–12. doi:10.1038/srep22989

17. Partridge JF, Borgstrøm B, Allshire RC. Distinct protein interaction domains and protein spreading in a complex centromere. Genes Dev. 2000;14: 783–791. doi:10.1101/gad.14.7.783

18. Volpe TA, Kidner C, Hall IM, Teng G, Grewal SIS, Martienssen RA. Regulation of heterochromatic silencing and histone H3 Lysine-9 methylation by RNAi. Science. 2002;297: 1833–1837. doi:10.1126/science.1074973

19. Folco HD, Pidoux AL, Urano T, Allshire RC. Heterochromatin and RNAi are required to establish CENP-A chromatin at centromeres. Science. 2008;319: 94–7. doi:10.1126/science.1150944

20. Janbon G, Ormerod KL, Paulet D, Byrnes EJ, Yadav V, Chatterjee G, et al. Analysis of the genome and transcriptome of Cryptococcus neoformans var. grubii reveals complex RNA expression and microevolution leading to virulence attenuation. PLOS Genet. 2014;10: e1004261. doi:10.1371/journal.pgen.1004261

21. Yadav V, Sun S, Billmyre RB, Thimmappa BC, Shea T, Lintner R, et al. RNAi is a critical determinant of centromere evolution in closely related fungi. Proc Natl Acad Sci. 2018; doi:10.1073/pnas.1713725115

22. Wang X, Hsueh Y-P, Li W, Floyd A, Skalsky R, Heitman J. Sex-induced silencing defends the genome of Cryptococcus neoformans via RNAi. Genes Dev. 2010;24: 2566–2582. doi:10.1101/gad.1970910

23. Wang X, Wang P, Sun S, Darwiche S, Idnurm A, Heitman J. Transgene induced co-suppression during vegetative growth in Cryptococcus neoformans. PLOS Genet. 2012;8: e1002885. doi:10.1371/journal.pgen.1002885

24. Janbon G, Maeng S, Yang D-H, Ko Y-J, Jung K-W, Moyrand F, et al. Characterizing the role of RNA silencing components in Cryptococcus neoformans. Fungal Genet Biol. 2010;47: 10. doi:10.1016/j.fgb.2010.10.005

25. Dumesic PA, Homer CM, Moresco JJ, Pack LR, Shanle EK, Coyle SM, et al. Product binding enforces the genomic specificity of a yeast polycomb repressive complex. Cell. 2015;160: 204–218. doi:10.1016/j.cell.2014.11.039

26. Catania S, Dumesic PA, Pimentel H, Nasif A, Stoddard CI, Burke JE, et al. Evolutionary persistence of DNA methylation for millions of years after ancient loss of a de novo methyltransferase. Cell. 2020;180: 263-277.e20. doi:10.1016/j.cell.2019.12.012

27. Feretzaki M, Billmyre RB, Clancey SA, Wang X, Heitman J. Gene network polymorphism illuminates loss and retention of novel RNAi silencing components in the Cryptococcus pathogenic species complex. PLOS Genet. 2016;12: e1005868. doi:10.1371/journal.pgen.1005868

28. Hagen F, Lumbsch HT, Arsic Arsenijevic V, Badali H, Bertout S, Billmyre RB, et al. Importance of resolving fungal nomenclature: the case of multiple pathogenic species in the Cryptococcus genus. mSphere. 2017;2. doi:10.1128/msphere.00238-17

29. Farrer RA, Desjardins CA, Sakthikumar S, Gujja S, Saif S, Zeng Q, et al. Genome evolution and innovation across the four major lineages of Cryptococcus gattii. mBio. 2015;6: e00868–15. doi:10.1128/mBio.00868-15

30. Scott KC, Sullivan BA. Neocentromeres: a place for everything and everything in its place. Trends Genet. 2013;30: 66–74. doi:10.1016/j.tig.2013.11.003

31. Nishimura K, Komiya M, Hori T, Itoh T, Fukagawa T. 3D genomic architecture reveals that neocentromeres associate with heterochromatin regions. J Cell Biol. 2019;218: 134–149. doi:10.1083/jcb.201805003

32. Shang W-HH, Hori T, Martins Nmcc, Toyoda A, Misu S, Monma N, et al. Chromosome engineering allows the efficient isolation of vertebrate neocentromeres. Dev Cell. Elsevier; 2013;24: 635–648. doi:10.1016/j.devcel.2013.02.009

33. Thakur J, Sanyal K. Efficient neocentromere formation is suppressed by gene conversion to maintain centromere function at native physical chromosomal loci in Candida albicans. Genome Res. 2013; 638–652. doi:10.1101/gr.141614.112

34. Ketel C, Wang HSW, McClellan M, Bouchonville K, Selmecki A, Lahav T, et al. Neocentromeres form efficiently at multiple possible loci in Candida albicans. PLOS Genet. 2009;5: e1000400. doi:10.1371/journal.pgen.1000400

35. Burrack LS, Hutton HF, Matter KJ, Clancey SA, Liachko I, Plemmons AE, et al. Neocentromeres provide chromosome segregation accuracy and centromere clustering to multiple loci along a Candida albicans chromosome. PLOS Genet. 2016;12: e1006317. doi:10.1371/journal.pgen.1006317

36. Scott KC, Bloom KS. Lessons learned from counting molecules: How to lure CENP-A into the kinetochore. Open Biol. 2014;4. doi:10.1098/rsob.140191

37. Sreekumar L, Jaitly P, Chen Y, Thimmappa BC, Sanyal A, Sanyal K. Cis-and trans-chromosomal interactions define pericentric boundaries in the absence of conventional heterochromatin. Genetics. 2019;212: 1121–1132. doi:10.1534/genetics.119.302179

38. Ishii K, Ogiyama Y, Chikashige Y, Soejima S, Masuda F, Kakuma T, et al. Heterochromatin integrity affects chromosome reorganization after centromere dysfunction. Science. 2008;321: 1088–91. doi:10.1126/science.1158699

39. Ohno Y, Ogiyama Y, Kubota Y, Kubo T, Ishii K. Acentric chromosome ends are prone to fusion with functional chromosome ends through a homology-directed rearrangement. Nucleic Acids Res. 2016;44: 232–244. doi:10.1093/nar/gkv997

40. Schotanus K, Heitman J. Centromere deletion in Cryptococcus deuterogattii leads to neocentromere formation and chromosome fusions. eLife. 2020;9: e56026. doi:10.7554/eLife.56026

41. Gröhs Ferrareze PA, Maufrais C, Streit RSA, Priest SJ, Cuomo C, Heitman J, et al. Application of an optimized annotation pipeline to the Cryptococcus deuterogattii genome reveals dynamic primary metabolic gene clusters and genomic impact of RNAi loss. bioRxiv. 2020; doi:https://doi.org/10.1101/2020.09.01.278374

42. Gent JI, Wang N, Dawe RK. Stable centromere positioning in diverse sequence contexts of complex and satellite centromeres of maize and wild relatives. Genome Biol. Genome Biology; 2017;18: 1–11. doi:10.1186/s13059-017-1249-4

43. Hori T, Kagawa N, Toyoda A, Fujiyama A, Misu S, Monma N, et al. Constitutive centromere-associated network controls centromere drift in vertebrate cells. J Cell Biol. 2017;216: 101–113. doi:10.1083/jcb.201605001

44. Smith KM, Phatale PA, Sullivan CM, Pomraning KR, Freitag M. Heterochromatin is required for normal distribution of Neurospora crassa CenH3. Mol Cell Biol. 2011;31: 2528–2542. doi:10.1128/MCB.01285-10

45. Wiemann P, Sieber CMK, von Bargen KW, Studt L, Niehaus EM, Espino JJ, et al. Deciphering the cryptic genome: genome-wide analyses of the rice pathogen Fusarium fujikuroi reveal complex regulation of secondary metabolism and novel metabolites. PLOS Pathog. 2013;9. doi:10.1371/journal.ppat.1003475

46. Fang Y, Coelho MA, Shu H, Schotanus K, Thimmappa, Bhagya C, Yadav V, et al. Long transposon-rich centromeres in an oomycete reveal divergence of centromere features in Stramenopila-Alveolata-Rhizaria lineages. PLOS Genet. 2019;16: e1008646. doi:doi.org/10.1371/journal.pgen.1008646

47. Schotanus K, Soyer JL, Connolly LR, Grandaubert J, Happel P, Smith KM, et al. Histone modifications rather than the novel regional centromeres of Zymoseptoria tritici distinguish core and accessory chromosomes. Epigenetics Chromatin. 2015;8: 41. doi:10.1186/s13072-015-0033-5

48. Mandáková T, Hloušková P, Koch MA, Lysak MA. Genome evolution in Arabideae was marked by frequent centromere repositioning. Plant Cell. 2020; doi:10.1105/tpc.19.00557

49. Henikoff S, Ahmad K, Malik HS. The centromere paradox: stable inheritance with rapidly evolving DNA. Science. 2001;293: 1098–102. doi:10.1126/science.1062939

50. Fukagawa T, Earnshaw WC. The centromere: Chromatin foundation for the kinetochore machinery. Dev Cell. 2014;30: 496–508. doi:10.1016/j.devcel.2014.08.016

51. Han F, Gao Z, Birchler JA. Reactivation of an inactive centromere reveals epigenetic and structural components for centromere specification in maize. Plant Cell. 2009;21: 1929– 1939. doi:10.1105/tpc.109.066662

52. Sato H, Masuda F, Takayama Y, Takahashi K, Saitoh S. Epigenetic inactivation and subsequent heterochromatinization of a centromere stabilize dicentric chromosomes. Curr Biol. 2012;22: 658–667. doi:https://doi.org/10.1016/j.cub.2012.02.062

53. Stimpson KM, Song IY, Jauch A, Holtgreve-Grez H, Hayden KE, Bridger JM, et al. Telomere disruption results in non-random formation of de novo dicentric chromosomes involving acrocentric human chromosomes. PLoS Genet. 2010;6: e1001061. doi:10.1371/journal.pgen.1001061

54. Alonso A, Hasson D, Cheung F, Warburton PE. A paucity of heterochromatin at functional human neocentromeres. Epigenetics and Chromatin. 2010;3: 6. doi:10.1186/1756-8935-3-6

55. Allshire RC, Javerzat JP, Redhead NJ, Cranston G. Position effect variegation at fission yeast centromeres. Cell. 1994;76: 157–169. doi:10.1016/0092-8674(94)90180-5

56. Castillo AG, Mellone BG, Partridge JF, Richardson W, Hamilton GL, Allshire RC, et al. Plasticity of fission yeast CENP-A chromatin driven by relative levels of histone H3 and H4. PLOS Genet. 2007;3: e121. doi:10.1371/journal.pgen.0030121

57. Nergadze SG, Piras FM, Gamba R, Corbo M, Cerutti F, McCarter JGW, et al. Birth, evolution, and transmission of satellite-free mammalian centromeric domains. Genome Res. 2018;28: 789–799. doi:10.1101/gr.231159.117

58. Purgato S, Belloni E, Piras FM, Zoli M, Badiale C, Cerutti F, et al. Centromere sliding on a mammalian chromosome. Chromosoma. 2015;124: 277–287. doi:10.1007/s00412-014-0493-6

59. Davidson RC, Blankenship JR, Kraus PR, D. M, Berrios J, Hull CM, et al. A PCR-based strategy to generate integrative targeting alleles with large regions of homology. Microbiology. 2002;2: 2607–2615.

60. Billmyre RB, Clancey SA, Heitman J. Natural mismatch repair mutations mediate phenotypic diversity and drug resistance in Cryptococcus deuterogattii. eLife. 2017;6: e28802. doi:10.7554/eLife.28802

61. Soyer JL, Möller M, Schotanus K, Connolly LR, Galazka JM, Freitag M, et al. Chromatin analyses of Zymoseptoria tritici : Methods for chromatin immunoprecipitation followed by high-throughput sequencing (ChIP-seq). Fungal Genet Biol. 2015;79: 63–70. doi:10.1016/j.fgb.2015.03.006

62. Langmead B. Aligning short sequencing reads with Bowtie. Curr Protoc Bioinforma. 2010; 11.7.1-11.7.14. doi:10.1002/0471250953.bi1107s32

63. Li H, Handsaker B, Wysoker A, Fennell T, Ruan J, Homer N, et al. The sequence alignment/map format and SAMtools. Bioinformatics. 2009;25: 2078–9. doi:10.1093/bioinformatics/btp352

64. Quinlan AR, Hall IM. BEDTools: A flexible suite of utilities for comparing genomic features. Bioinformatics. 2010;26: 841–842. doi:10.1093/bioinformatics/btq033

65. Ramírez F, Dündar F, Diehl S, Grüning BA, Manke T. DeepTools: A flexible platform for exploring deep-sequencing data. Nucleic Acids Res. 2014;42: 187–191. doi:10.1093/nar/gku365

66. Feng J, Liu T, Qin B, Zhang Y, Liu XS. Identifying ChIP-seq enrichment using MACS. Nat Protoc. Nature Publishing Group; 2012;7: 1728–1740. doi:10.1038/nprot.2012.101

67. Kim D, Pertea G, Trapnell C, Pimentel H, Kelley R, Salzberg SL. TopHat2: accurate alignment of transcriptomes in the presence of insertions, deletions and gene fusions. Genome Biol. BioMed Central Ltd; 2013;14: R36. doi:10.1186/gb-2013-14-4-r36

68. Schneider RO, de Souza Süffert Fogaça N, Kmetzsch L, Schrank A, Vainstein MH, Staats CC. Zap1 regulates zinc homeostasis and modulates virulence in Cryptococcus gattii. PLoS One. 2012;7: e43773. doi:10.1371/journal.pone.0043773

69. Priest SJ, Coelho MA, Mixão V, Clancey S, Xu Y, Sun S, et al. Factors enforcing the species boundary between the human pathogens Cryptococcus neoformans and Cryptococcus deneoformans. PLOS Genet. 2020;17: e1008871. doi:10.1371/journal.pgen.1008871

70. Findley K, Rodriguez-Carres M, Metin B, Kroiss J, Fonseca A, Vilgalys R, et al. Phylogeny and phenotypic characterization of pathogenic Cryptococcus species and closely related saprobic taxa in the tremellales. Eukaryot Cell. 2009;8: 353–361. doi:10.1128/EC.00373-08

71. Yadav V, Sun S, Coelho MA, Heitman J. Centromere scission drives chromosome shuffling and reproductive isolation. Proc Natl Acad Sci U S A. 2020;117: 7917–7928. doi:10.1073/pnas.1918659117

72. Simpson JT, Workman RE, Zuzarte PC, David M, Dursi LJ, Timp W. Detecting DNA cytosine methylation using nanopore sequencing. Nat Methods. Nature Publishing Group; 2017;14: 407–410. doi:10.1038/nmeth.4184

73. Pitkin JW, Panaccione DG, Walton JD. A putative cyclic peptide efflux pump encoded by the TOXA gene of the plant-pathogenic fungus Cochliobolus carbonum. Microbiology. 1996;142: 1557–1565. doi:10.1099/13500872-142-6-1557

